# Inverse expression of Ten3 and Lphn2 across the developing mouse brain suggests a global strategy for circuit assembly

**DOI:** 10.1101/2025.08.13.670004

**Authors:** URee Chon, Daniel T. Pederick, Jun Ho Song, Yanbo Zhang, Isabella Rana, Liqun Luo

**Author notes:** These authors contributed equally to this study (URee Chon, Daniel T. Pederick). Department of Neuroscience, Johns Hopkins University School of Medicine, Baltimore, MD 21205, USA. Neuroscience PhD Program, Columbia University, New York, NY 10032, USA.

## Abstract

Precise wiring of neural circuits requires molecular strategies that ensure accurate target selection across diverse brain regions. Here, we identify inverse expression between a ligand–receptor pair, Teneurin-3 (Ten3) and Latrophilin-2 (Lphn2), across the developing mouse brain. Ten3 and Lphn2 exhibit inverse expression gradients along a retinotopic axis orthogonal to the ephrin-A and EphA gradients; along the tonotopic axis across multiple brainstem auditory nuclei; and along the dorsomedial–ventrolateral axis in striatum and pallidum. Their inverse expression also creates discrete domains of cerebellar Purkinje cells and cerebellar nuclei. Using conditional tag mice, we show that inverse Ten3 and Lphn2 expression patterns predict connectivity, following a ‘Ten3→Ten3, Lphn2→Lphn2’ rule in all above circuits, and that Lphn2 is required in executing this rule in Purkinje cells→cerebellar nuclei projection. Our findings suggest a global strategy of coordinating gene expression of key wiring molecules with circuit connectivity across the developing brain.

## INTRODUCTION

Precise wiring of neural circuits during development is essential for proper information processing and animal behavior. The specificity of neuronal connectivity emerges from cell–cell interactions that rely on the differential expression of cell surface proteins (CSPs) to distinguish neuronal identity and guide the correct selection of synaptic partners.^1–5^ However, while the human brain contains ∼10¹¹ neurons that form connections, the genome encodes only ∼20,000 protein-coding genes, less than ∼3,000 of which encode CSPs.^6,7^ This disparity raises a fundamental question: how does the brain achieve such extraordinary wiring specificity with a limited molecular repertoire? Several mechanisms have been proposed to reconcile this disparity^4^, such as graded expression to enable different levels of the same CSP to specify different connections^8,9^ and combinatorial actions of multiple CSPs to match synaptic partners.^10^ Despite extensive research on individual or pairs of CSPs that act as guidance molecules for wiring of specific circuits, however, the spatial dynamics of ligand–receptor pairs and how they cooperate to establish precise wiring of multiple circuits across development have rarely been examined.

Teneurins are evolutionarily conserved type II transmembrane protein that instructs synaptic partner matching by homophilic adhesion.^11–14^ Additionally, Teneurin-3 (Ten3) and the adhesion G-protein-coupled receptor Latrophilin-2 (Lphn2) display inverse expression in two parallel hippocampal networks.^15^ Ten3^+^ proximal CA1 axons connect with Ten3^+^ target neurons in distal subiculum, and Lphn2^+^ distal CA1 axons connect with Lphn2^+^ target neurons in proximal subiculum. Ten3 and Lphn2 cooperate for CA1→subiculum target through Ten3–Ten3 homophilic attraction and Ten3–Lphn2 reciprocal heterophilic repulsion.^15,16^ In a companion manuscript, we demonstrate that the Ten3–Lphn2 molecular module is repeatedly used to instruct target selection across multiple nodes of the extended hippocampal network.^17^ These studies raise the question of whether this ligand–receptor pair could be broadly deployed in establishing wiring specificity in other regions of the mammalian brain.

Here we report, using whole-brain expression at multiple developmental stages, that inverse expression of Ten3 and Lphn2 is present in many regions across the developing mouse brain. We focus our detailed analyses on four systems. These include the visual and auditory systems, in which nearby neurons in the input field project axons topographically to nearby neurons in the target field to form continuous retinotopic and tonotopic maps^18,19^, as well as the basal ganglia and cerebellum where the connectivity organization is less well defined. Across these regions, we observe that inverse Ten3 and Lphn2 expression patterns predict wiring specificity, following a ‘Ten3→Ten3, Lphn2→Lphn2’ rule relating CSP expression to connectivity. In the visual system, Ten3–Lphn2 inverse expression defines retinotopy along the dorsal–ventral axis orthogonal to the anterior–posterior axis defined by inverse ephrin-A–EphA expression.^18,20^ In the auditory system, Ten3–Lphn2 inverse expression aligns with tonotopy across multiple brainstem auditory nuclei, providing a logic for how information about sound frequency could be relayed faithfully across multiple synapses. In the basal ganglia, Ten3–Lphn2 inverse expression coincides with striatal-pallidal and cortical-striatal projections. In the cerebellum, Ten3–Lphn2 inverse expression matches Purkinje cell→cerebellar nuclei projection specificity, and we further demonstrate that Lphn2 is required for implementing the ‘Ten3→Ten3’ connectivity rule. These findings, along with our companion manuscripts^17,21^, establish Ten3 and Lphn2 as a molecular module that organizes specific and precise connectivity across anatomically and functionally distinct brain and spinal cord regions, uncovering a global principle for circuit organization throughout the developing nervous system.

## RESULTS AND DISCUSSION

### Ten3 and Lphn2 exhibit inverse expression across many regions of the developing brain

Given the inverse Ten3 and Lphn2 expression in the extended hippocampus network,^15,17^ we first asked whether similar patterns extend to other regions of the brain during development. We utilized knock-in alleles of Ten3-HA (**Figure S1A**) and Lphn2-mVenus^22^ to characterize their distribution across the entire developing brain at embryonic day (E) 15 and 17, and postnatal day (P) 0, 2, 4, and 8. Immunostained brains were cleared and whole-mount imaged with light-sheet microscopy (**Figure 1A**).

**Figure 1.**
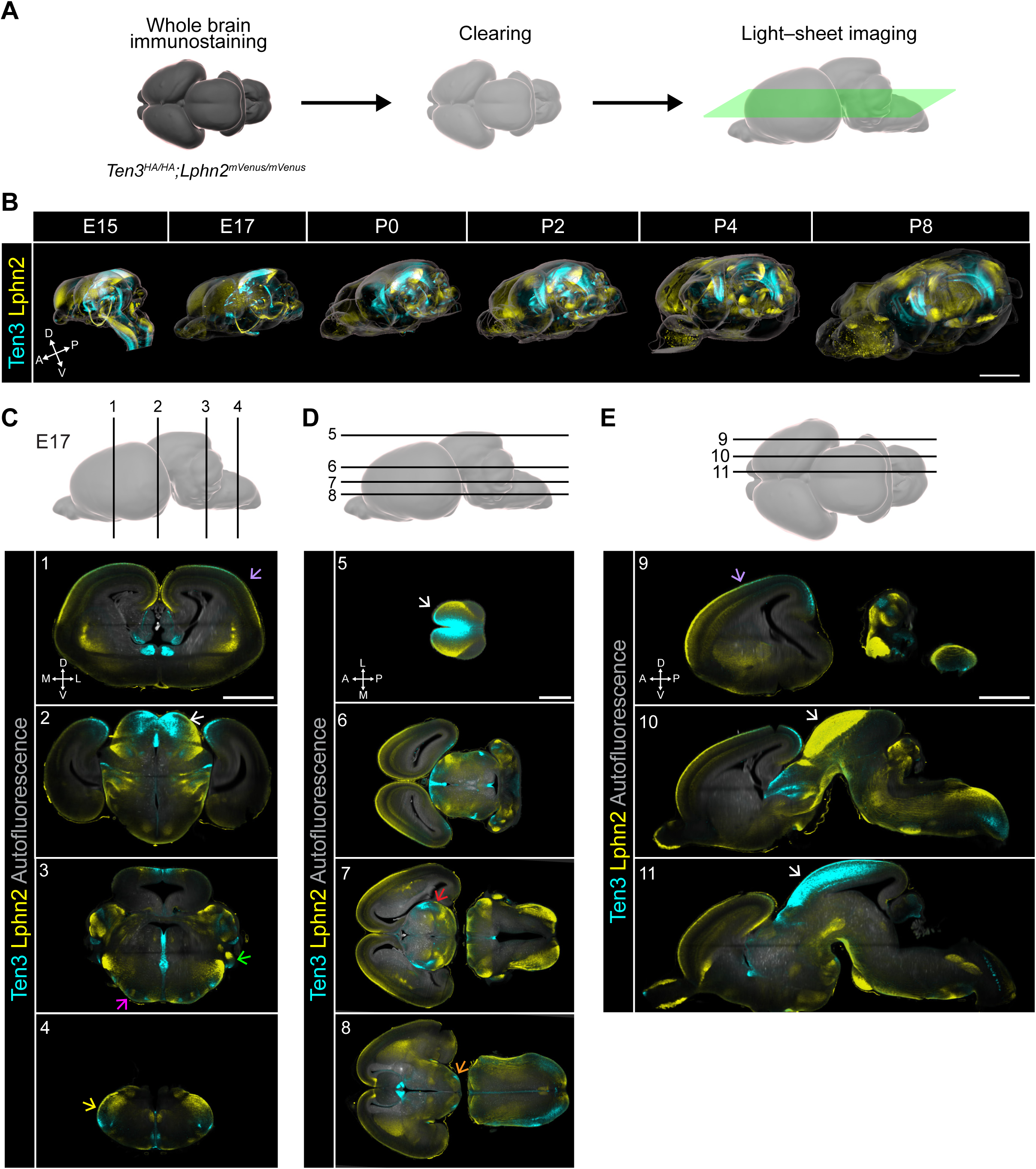
Whole brain staining of Ten3 and Lphn2 across development. (A) Overview of the procedure used to stain whole mouse brains for Ten3 and Lphn2 proteins. (B) 3D staining patterns of Ten3 and Lphn2 proteins in the mouse brain across different stages of development. E, embryonic day; P, postnatal day. Scale bar, 2 mm. (C–E) Ten3 and Lphn2 protein expression in single optical sections in the coronal (C), horizontal (D), and sagittal (E) planes of the same 3D volume of an E17 brain imaged by light sheet microscopy. The number of each image is matched with the section location (line) in the schematic above. Superior colliculus, white arrows; superior olivary complex, magenta arrow; ventral cochlear nucleus, green arrow; spinal trigeminal nucleus, yellow arrow; thalamic nuclei, red arrow; mammillary nucleus of the hypothalamus, orange arrow; neocortex, purple arrows. Detailed expression patterns in brain regions of E17 and P2 mouse brains can be found in **Extended Data 1**. Scale bars, 1 mm. In this and all subsequent figures, A, anterior; P, posterior; D, dorsal; V, ventral; M, medial; L, lateral. Unless otherwise mentioned, all Ten3 and Lphn2 proteins were visualized with anti-HA and anti-GFP antibodies in *Ten3^HA/HA^*;*Lphn2^mV/mV^*(mV for mVenus) knock-in mice and shown as cyan and yellow colors, respectively. See Figure S1 and Videos S1 and S2 for additional data.

Across development, we observed Ten3 and Lphn2 expression in broad, largely non-overlapping and often adjacent regions spanning the forebrain, midbrain, and hindbrain (**Figure 1B**; **Videos S1, S2**). For example, at E17, we observed differential expression in discrete brain regions, including the neocortex, striatum, hypothalamus, superior colliculus, cerebellar cortex, cochlear nucleus, spinal nucleus of the trigeminal, and nucleus of the solitary tract through coronal (**Figure 1C**), horizontal (**Figure 1D**), and sagittal (**Figure 1E**) sections. As examples, Ten3 was enriched medially and Lphn2 laterally in the superior colliculus (**Figure 1C_2_, D_5_, E_10_, E_11_**, white arrows). Conversely, Lphn2 was enriched medially and Ten3 laterally in multiple nuclei of the superior olivary complex (**Figure 1C_3_**, magenta arrow), as well as the mammillary nucleus of the hypothalamus (**Figure 1D_8_**, orange arrow). In the cochlear nucleus (**Figure 1C_3_**, green arrow) and spinal trigeminal nucleus (**Figure 1C_4_**, yellow arrow), Lphn2 expression was higher in dorsal and Ten3 higher in ventral regions. Ten3 and Lphn2 also exhibited inverse expression in discrete nuclei of the thalamus (**Figure 1D_7_**, red arrow). In the neocortex, Ten3 was high posteriorly whereas Lphn2 was high anteriorly (**Figure 1E_9_**, purple arrow); at a coronal section midway along the anterior–posterior axis, Lphn2 was high at lateral and medial edges whereas Ten3 was high in the intermediate zone (**Figure 1C_1_**, purple arrow). To facilitate the identification of anatomical regions that express Ten3 and Lphn2 throughout the entire brain and across different timepoints, we developed an interactive, ImageJ-based visualization tool that enables users to actively navigate and examine anatomical regions of interest (**Extended Data 1**).

Together, these data show that Ten3 and Lphn2 proteins exhibit spatially segregated and largely inverse expression patterns across the developing brain. These inverse expression patterns do not generally follow fixed body axes but rather are specialized in each brain region. We focused further investigation on four systems that exhibited robust and stereotyped Ten3–Lphn2 inverse expression: the visual system, auditory system, basal ganglia, and cerebellum, in the context of development and connectivity patterns of each system. Quantitative protein level analyses across multiple developmental stages examined revealed dynamic expression of Ten3 and Lphn2 across developmental stages, and that in systems where timing of target selection of axons has been determined, this timing overlaps with peak expression of Ten3 and Lphn2 (**Figure S2**).

### Ten3 and Lphn2 expression aligns with retinotopy along the dorsal–ventral axis of the retina

Retinotopy is a fundamental organization of the visual system—retinal ganglion cells (RGCs) in nearby positions of the retina project axons to nearby neurons in the target regions including the superior colliculus (SC), such that the two-dimensional spatial map of the world in the retina is recapitulated in brain targets (**Figure 2A**). The orderly projection of RGC axons to the SC (tectum in non-mammalian vertebrates) have been well described: RGCs from the anterior (nasal) retina project to posterior SC/tectum, posterior (temporal) retina to anterior SC/tectum, ventral retina to medial SC/tectum, and dorsal retina to lateral SC/tectum.^1,20^ An extensive body of work has demonstrated that graded expression of cell-surface proteins ephrin-As and their receptors EphAs in both the retina and SC instructs retinotopic map along the anterior–posterior axis of the retina.^8,9,23–26^ Much less is known about the mechanisms that control RGC axon targeting along the orthogonal dorsal–ventral axis, even though several candidates have been proposed, including ephrin-B/EphB and the Wnt pathway. Ephrin-Bs and EphBs appear to play different roles in frogs and mice, exhibit modest differential expression, and produce mild phenotypes with *EphB* disruption;^27,28^ Wnt signaling lacks loss-of-function evidence to support a direct role.^29^ Interestingly, *Ten3* mRNA has been reported to be expressed in the ventral retina and was shown to have defects in retinocollicular circuit wiring when deleted.^30^ These findings suggested Ten3 as a candidate guidance molecule that may contribute to dorsal–ventral retinotopic organization.

**Figure 2.**
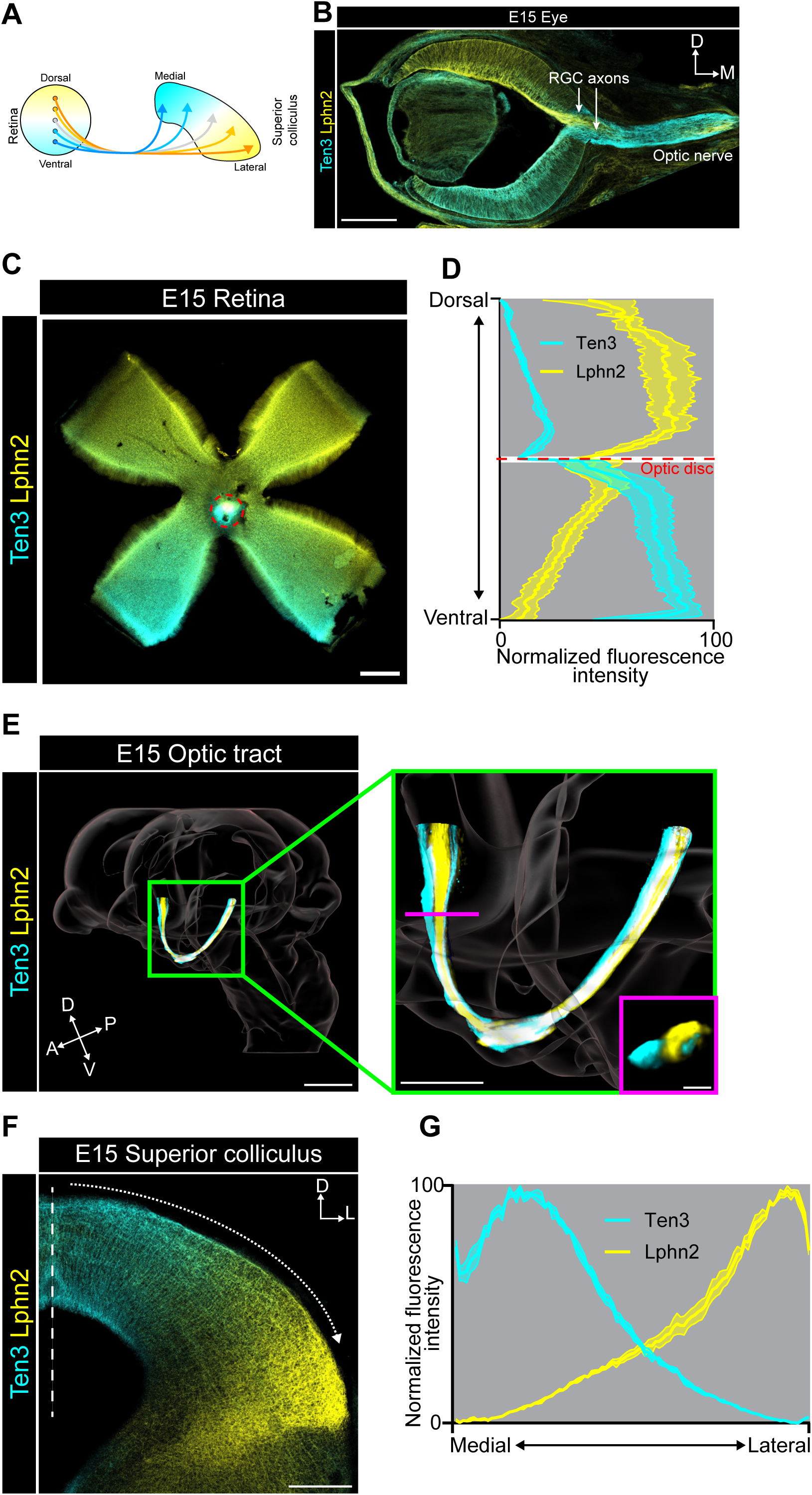
Inverse expression patterns of Ten3 and Lphn2 follow retinotopic connections. (A) Schematic diagram of retinotopic connections from the retina to the superior colliculus along the dorsal–ventral axis of the retina, with Ten3 (cyan) and Lphn2 (yellow) expression from our data superimposed. (B) Ten3 and Lphn2 protein expression in a single coronal section of the retina. RGC axons, retinal ganglion cell axons. Scale bar, 200 µm. (C) Flat whole mount E15 retina preparation stained for Ten3 and Lphn2 proteins. Red circle indicates optic nerve head, where RGC axons exit the retina. Scale bar, 200 µm. (D) Quantification of Ten3 and Lphn2 proteins across the dorsal–ventral axis of the retina. See STAR Methods for details on regions analyzed (n = 4 mice; 1 retina/mouse). Mean ± SEM. (E) 3D reconstruction of E15 optic tract from light sheet microscope images with Ten3 and Lphn2 proteins labeled. Scale bar, 400 µm. Green inset, zoom in of the optic tract (scale bar, 200 µm). Magenta inset, cross section of the optic tract (scale bar, 50 µm). Both insets reveal two discrete Ten3^+^ bundles in the most anterior and posterior regions of the optic tract, along with a single Lphn2^+^ bundle in the middle. (F) Ten3 and Lphn2 protein expression in the superior colliculus in a coronal section of an E15 brain. Vertical dashed line indicates the midline. Dotted arrow indicates the axis used for quantification in (G). Scale bar, 200 µm. (G) Quantification of Ten3 and Lphn2 protein across the medial–lateral axis of the superior colliculus (n = 3 mice). Mean ± SEM. See Figures S2 and S3 for additional data.

RGC projections and target selection at the SC begin during embryogenesis (**Figure S2A**).^31^ Using immunostaining in a coronal section at E15, we found that Ten3 and Lphn2 was highly expressed in the ventral and dorsal retina, respectively (**Figure 2B**). Quantification of normalized fluorescence intensity from flat-mount retina further revealed that Ten3 and Lphn2 proteins formed inverse gradients along the dorsal–ventral axis of the retina (**Figure 2C, D**). Three-dimensional reconstructions of cleared E15 whole brains visualized by light-sheet microscopy revealed that Ten3 and Lphn2 labeled discrete axon bundles in the optic tract, which comprises RGC axons connecting the eyes to brain targets (**Figure 2E**). Inverse expression was also evident in the SC, with preferential expression of Ten3 in medial SC, inversely to Lphn2 enrichment in lateral SC (**Figure 2F**; **Figure S2A**), forming opposing gradients (**Figure 2G**). Double *in situ* hybridization for *Ten3* and *Lphn2* mRNA confirmed local production of each CSP’s mRNA in the SC (**Figure S3**), suggesting that the presence of Ten3 and Lphn2 protein in SC is likely contributed by both RGC axons and SC target neurons. Given the known topographic projections of ventral and dorsal RGCs target medial and lateral SC,^1^ our findings indicate that RGC axon projections follow the ‘Ten3→Ten3, Lphn2→Lphn2’ rule, and suggest that these proteins could participate in establishing the retinocollicular maps along the dorsal–ventral axis, perhaps in collaboration ephrin-B/EphB and Wnt,^27–29^ orthogonal to the ephrin-A/EphA-regulated anterior–posterior axis.

### Ten3 and Lphn2 expression aligns with tonotopy in multiple brainstem nuclei along the ascending auditory pathway

Tonotopy is a fundamental organizing principle of the auditory system.^32^ Sounds of different frequencies are represented by hair cells at different locations in the cochlea. Through ordered axonal projections, this tonotopic organization is recapitulated along the central auditory pathway in brainstem nuclei (**Figure 3A**) and beyond. These tonotopic maps are fundamental to frequency discrimination, pitch perception, and language processing.^33,34^ Little is known about the molecular mechanisms that instruct the formation of tonotopic connections.^19^

**Figure 3.**
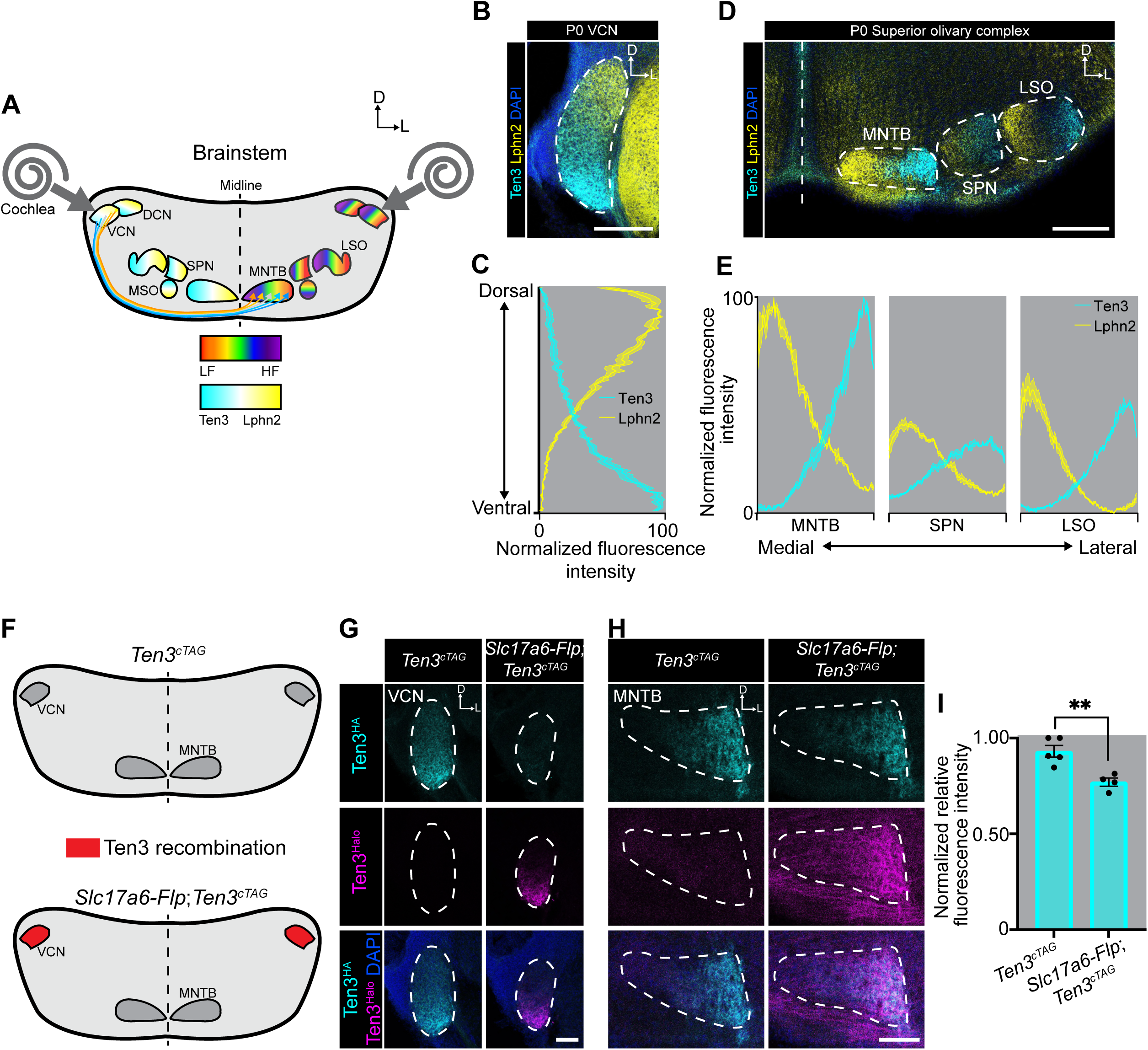
Tonotopic expression of Ten3 and Lphn2 in auditory processing regions of the brainstem. (A) Schematic diagram of auditory processing nuclei in the brainstem and tonotopic organization represented in a rainbow color spectrum. Cyan and yellow summarize expression patterns of Ten3 and Lphn2 from our data, which align with tonotopic maps across these nuclei. Tonotopic projections from VCN to MNTB are illustrated. DCN, dorsal cochlear nucleus; VCN, ventral cochlear nucleus; MSO, medial superior olive; LSO, lateral superior olive; SPN, superior paraolivary nucleus; MNTB, medial nucleus of the trapezoid body. (B) Ten3 and Lphn2 protein expression in the VCN of a P0 brain. Scale bar, 200 µm. (C) Quantification of Ten3 and Lphn2 protein across the dorsal–ventral axis of the VCN (n = 3 mice). Mean ± SEM. (D) Ten3 and Lphn2 protein expression in the superior olivary complex of a P0 brain. Scale bar, 200 µm. (E) Quantification of Ten3 and Lphn2 protein across the medial–lateral axis of the superior olivary complex (n = 3 mice). Mean ± SEM. (F) Schematic diagram depicting spatial patterns of recombination of *Ten3^cTAG^* allele in the VCN→MNTB projection pathway in the absence (top) and presence (bottom) of *Slc17a6-Flp*. (G, H) Ten3^HA^ and Ten3^Halo^ protein expression in the VCN (G) and MNTB (H) of *Ten3^cTAG/cTAG^* and *Slc17a6-Flp;Ten3^cTAG/cTAG^* P0 mice. VCN (G) and MNTB (H) are dash outlined. Scale bar, 100 µm. (I) Quantification of relative HA staining in the MNTB of *Ten3^cTAG/cTAG^* (n = 5) and *Slc17a6-Flp;Ten3^cTAG/cTAG^* (n = 4) P0 mice. **, p < 0.01; unpaired t-test. See Figures S1, S2, S4, and S5 for additional data.

Immunostaining revealed that during embryonic and early postnatal periods, Ten3 and Lphn2 proteins were distributed in opposing gradients in all cochlear nuclei including the ventral cochlear nucleus (VCN; **Figure 3B, C**; **Figure S2B**), as well as medial nucleus of the trapezoid body (MNTB), superior paraolivary nucleus (SPN), and lateral superior olive (LSO) in the superior olivary complex (**Figure 3D, E**; **Figure S2C**). Strikingly, Ten3 expression always aligned with low-frequency regions, whereas Lphn2 with high-frequency regions (compare **Figure 3B–E** with **Figure 3A**). Double *in situ* hybridization for *Ten3* and *Lphn2* mRNA confirmed local production of Ten3 and Lphn2 in all nuclei of the auditory brainstem (**Figure S4**), suggesting that the protein expression is likely contributed by both input axons and target neurons. Given the known topographic projections between low- and high-frequency regions, these expression patterns indicate that the projections of auditory neurons between different nuclei follow the ‘Ten3→Ten3, Lphn2→Lphn2’ connectivity rule, and suggest that these proteins could participate in establishing tonotopic maps prior to the onset of hearing.

### Analyzing Ten3 expression separately in axons and targets in the VCN→MNTB projections

The ‘Ten3→Ten3, Lphn2→Lphn2’ connectivity rule implies that in each brain region that follows this rule, Ten3 or Lphn2 proteins are contributed by both input axons and target neurons. To directly visualize this and to quantify the contributions of proteins produced by axons and targets, we examined the contribution of Ten3 in axons and targets separately, using the VCN→MNTB projections as an example. To differentially label the same protein in axons and targets, we generated conditional tag (*cTAG*) mice for Ten3 (**Figure S1A**). In *Ten3^cTAG^* mice, the endogenous Ten3 is tagged with an HA epitope tag, which is converted to a Halo-tag in the presence of Flp recombinase. Expression analysis in the hippocampus before and after introducing a germline-FlpO validated the fidelity of protein expression as previously reported^13,15^ (**Figure S1A**).

Projection neurons from the VCN to the MNTB are excitatory neurons that express *Slc17a6* encoding the vesicular glutamate transporter VGLUT2, whereas neurons in the MNTB are exclusively inhibitory.^35,36^ To selectively label Ten3^+^ excitatory projection neurons, we crossed *Ten3^cTAG^* mice with *Slc17a6-Flp*, which expresses Flp in all VGLUT2^+^ glutamatergic neurons, including those in the VCN (**Figure S5B**). This strategy enabled us to independently examine Ten3^Halo^ from VCN axons and Ten3^HA^ produced by local MNTB neurons at the MNTB (**Figure 3F**).

In control *Ten3^cTAG^* mice lacking Flp expression, Ten3^HA^ (but no Ten3^Halo^) was detected in the VCN, consistent with the absence of recombination. In *Slc17a6-Flp;Ten3^cTAG^* mice, we observed robust recombination throughout VCN, with Ten3^Halo^ completely replacing Ten3^HA^ by P0 (**Figure 3G**). At the target MNTB, *Ten3^cTAG^* mice displayed Ten3^HA^ signal but no Ten3^Halo^ signal, consistent with lack of recombinase activity in axons and targets. By contrast, *Slc17a6-Flp;Ten3^cTAG^* mice exhibited Ten3^Halo^ signal that overlapped with Ten3^HA^ expression in the lateral region of the MNTB (**Figure 3H**). Thus, axons of Ten3^Halo+^ VCN excitatory neurons preferentially target to regions enriched for Ten3^HA+^ inhibitory neurons in the MNTB.

Our data reveal that Ten3 is predominantly expressed in excitatory neurons in the VCN, as in *Slc17a6-Flp;Ten3^cTAG^*mice, we did not detect any residual Ten3^HA^signals (**Figure 3G**, right panels). Moreover, given the complete conversion of Ten3^HA^ to Ten3^Halo^ in VCN, Ten3^HA^ signals in the MNTB are not contributed by VCN axons and derive exclusively from local MNTB neurons. This allowed us to quantify the relative contribution of Ten3 proteins in axons and targets. Comparing Ten3^HA^ levels in the MNTB in *Ten3^cTAG^* and *Slc17a6-Flp;Ten3^cTAG^*mice indicates that the majority of Ten3 protein in the MNTB was contributed by target neurons (**Figure 3I**). Finally, Ten3^Halo+^ VCN axons terminated at Ten3^HA^ region in the lateral MNTB, providing direct evidence supporting the ‘Ten3→Ten3’ targeting rule. Interestingly, Ten3^Halo+^ VCN axons passed by Ten3^−^(and Lphn2^+^) regions in the medial MNTB to reach their destination (**Figure 3H**, middle row). This observation highlights the selectivity of VCN axon targeting and suggests that repulsion from Lphn2 may contribute to this selectivity, as previously described in the hippocampal network.^15,17^

### Ten3 and Lphn2 display inverse expression in the basal ganglia

We next focused on the basal ganglia, a key regulator of motor output, action selection, and decision making.^37,38^ GABAergic spiny projection neurons in the input nucleus of the basal ganglia, the caudate putamen (CP, also known as striatum) form a direct pathway projecting to the globus pallidus internal segment (GPi) and substantia nigra, and an indirect pathway that first projects to the globus pallidus external segment (GPe). We found that Ten3 and Lphn2 exhibited inverse expression in both CP and GPe (**Figure S2D**; **Video S1**), and thus focused our further analysis on the CP→GPe pathway (**Figure 4A**).

**Figure 4.**
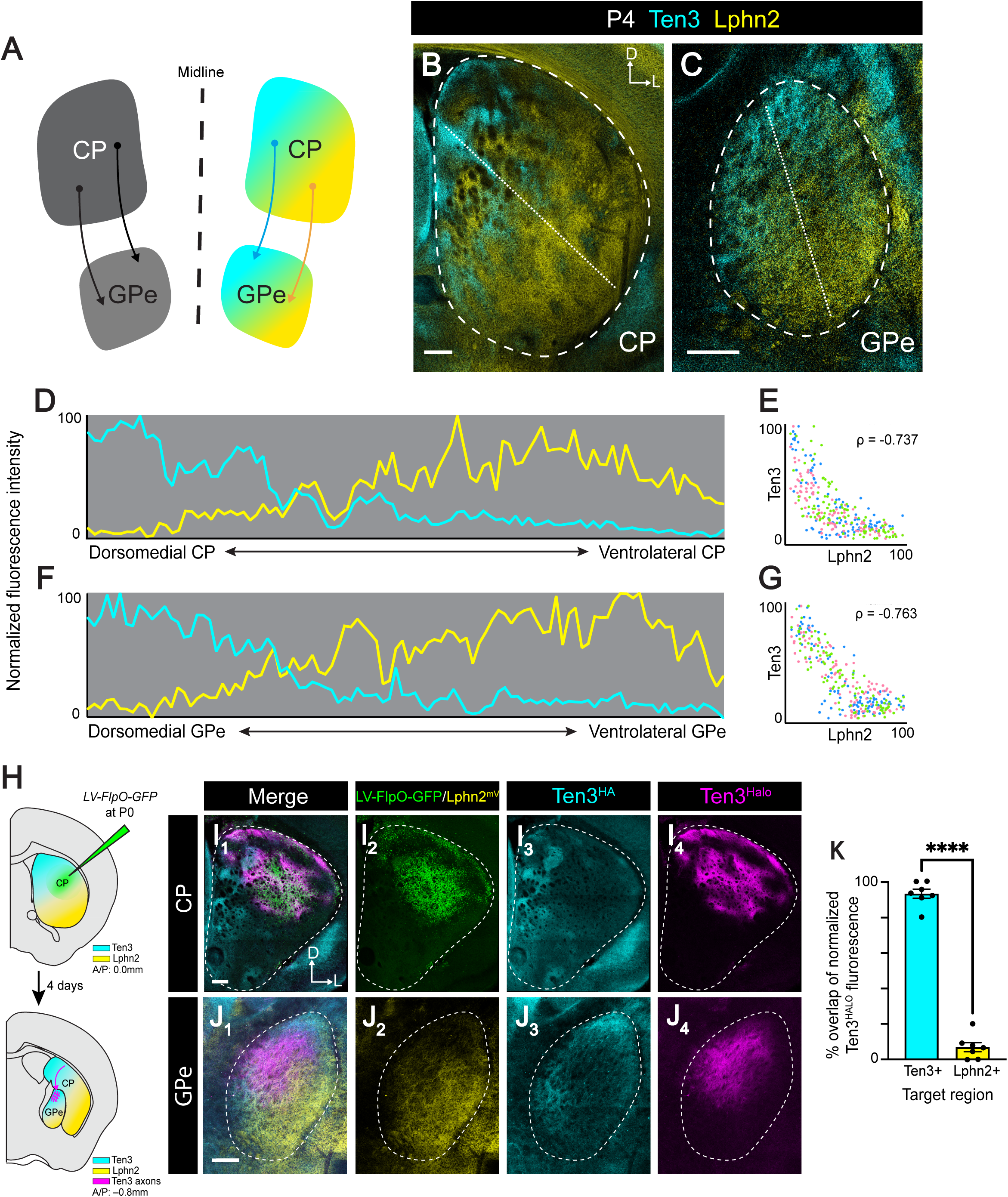
Ten3 and Lphn2 display inverse expression in the CP and GPe. (A) Schematic diagram of connections from the caudate putamen (CP) to the globus pallidus external segment (GPe), constituting part of the indirect pathway of the basal ganglia. (B, C) Ten3 and Lphn2 protein expression in the CP (B) and GPe (C) of a P4 brain, shown in coronal sections. CP and GPe are demarcated by the dashed outlines. Dotted lines indicate the axis used for quantification in (D) and (F). Scale bar, 200 µm. (D) Quantification of Ten3 and Lphn2 protein across the dorsomedial–ventrolateral axis (dotted line in B) of the CP denoted. (E) Spearman’s correlation between Ten3 and Lphn2 expression in the CP (n = 3 mice). Each dot represents the normalized fluorescence intensity of Ten3 and Lphn2 in a single x-y spatial bin along the dorsomedial–ventrolateral axis (n = 100 bins per brain). Dots of the same color are from the same mouse. See STAR Methods for details on analysis. Spearman’s correlation, ****p < 0.0001. (F, G) Same as (D, E), except for GPe. Spearman’s correlation, ****p < 0.0001. (H) Experimental design and summary of results for tracing CP→GPe connectivity pattern. LV, lentivirus. Recombination of *Ten3^cTAG^*allele at the injection site causes Ten3^HA+^ CP neurons to express Ten3^Halo^. Recombined Ten3^Halo+^ CP axons target GPe regions that express Ten3^HA^. (I) Injection site of *LV-FlpO-GFP* in the CP of *Ten3^cTAG/cTAG^;Lphn2 ^mV/mV^* mice at P4. *LV-FlpO-GFP* injected CP neurons shown in green, Ten3^HA^ in cyan, and Ten3^Halo^ in magenta. Dotted lines outline the CP. Scale bar, 200 µm. (J) Projection patterns of Ten3^Halo+^ CP axons (from injection in panel I) in the GPe. Ten3^Halo+^ CP axons target the Ten3^HA+^ region and avoid Lphn2^mV+^ region of the GPe. Dotted lines outline the GPe. Scale bar, 200 µm. (K) Percent overlap of normalized fluorescence of Ten3^Halo+^ CP projections to Ten3^HA+^ or Lphn2^mV+^ GPe target regions in *Ten3^cTAG/cTAG^;Lphn2 ^mV/mV^* mice at P4 (n = 7). Paired t-test; ****p < 0.0001. See Figures S2 and S6 and Video S3 for additional data.

Immunostaining of Ten3 and Lphn2 revealed preferential expression of Ten3 in dorsomedial CP and GPe globally, inversely to Lphn2 enrichment in ventrolateral CP and GPe (**Figure 4B, C**; **Figure S2D**).

Quantification across the dorsomedial–ventrolateral axis of the CP and GPe revealed inverse gradients of expression (**Figure 4D, F**). This inverse pattern was confirmed by Spearman’s correlation analysis across animals, revealing significant negative correlations between Ten3 and Lphn2 expression in both the CP (**Figure 4E**) and GPe (**Figure 4G**). Three-dimensional reconstructions of cleared whole brains visualized by light-sheet microscopy reinforced the consistent spatial segregation of Ten3 and Lphn2 in the CP and GPe (**Video S3**). Double *in situ* hybridization for *Ten3* and *Lphn2* mRNA showed that both *Ten3* and *Lphn2* mRNAs were expressed in the CP and GPe (**Figure S6A, B**). These experiments validated that at least a part of the proteins we visualized in both the CP and GPe were produced locally rather than exclusively contributed by incoming axons. Furthermore, within CP, both Ten3 and Lphn2 mRNAs were both co-expressed with *Tac1* or *Penk*, markers for spiny projection neurons in the direct and indirect pathways, respectively (**Figure S6C, D**).

Unlike the continuous gradients at embryonic stages (**Figure S2D**), Ten3^+^ regions in the dorsomedial CP formed spatially clustered hotspots by P4, intermingled with cortical axon bundles (**Figure 4B**). Lphn2^+^ regions maintained a broader distribution of expression throughout the ventrolateral CP. Near the central region of the CP where the global gradients of Ten3 and Lphn2 met, Lphn2^+^ regions also formed hotspots interdigitating with the Ten3^+^ hotspots (**Figure 4B**; **Figure S2D**). To test whether these hotspots are related to previously described striosome (patch) and matrix compartments,^39^ we performed co-expression of Ten3 and Lphn2 with μ-opioid receptor (MOR) or calbindin, markers of striosomes and matrix, respectively.^39,40^ We did not find clear correspondence between their expression patterns (**Figure S6E, F**), suggesting that heterogeneity created by Ten3 and Lphn2 hotspots is unrelated to striosomes and matrix.

### Striatopallidal projections follow the Ten3→Ten3 rule

Given that the CP and GPe both exhibit differential expression of Ten3 and Lphn2, we next asked whether the CP→GPe projections follow the ‘Ten3→Ten3’ rule. We injected lentivirus expressing *FlpO-GFP* into the CP of *Ten3^cTAG/cTAG^;Lphn2 ^mV/mV^* mice at P0 and examined projection patterns of Ten3^Halo+^ CP axons in the GPe at P4 (**Figure 4H**). The robust Ten3^Halo^ signal confirmed local production of Ten3 in the CP. We found that Ten3^Halo+^ axons from the CP preferentially innervate the Ten3^HA^ regions of the GPe, while avoiding Lphn2^mV^ regions (**Figure 4I, J**). Quantification of axon-target overlap confirmed substantial overlap of Ten3^Halo+^ CP axon signals within Ten3^HA^ target regions but little overlap in the Lphn2^mV^ target region (**Figure 4K**). Thus, the CP→GPe projections follow the ‘Ten3→Ten3’ rule.

### Corticostriatal projections follow the Ten3→Ten3 rule

The CP receives excitatory input from across the neocortex, as well as specific thalamic regions, mainly the parafascicular nuclei.^41^ We next examined whether these input axons contribute to Ten3 expression in the CP, and whether input projections also follow the ‘Ten3→Ten3’ rule, utilizing our conditional tag allele in combination with cell-type-specific Flp drivers to differentially label Ten3 produced from input axons versus the target arbors.

Before describing the contribution of thalamic and cortical inputs of Ten3 to the CP, we performed two control experiments. First, in the absence of Flp recombinase expression, we did not detect Ten3^Halo^ signal in the CP of *Ten3^cTAG^*mice (**Figure 5A**). Second, to visualize local Ten3 production in CP neurons, which are predominantly GABAergic, we crossed *Ten3^cTAG^* mice with *Slc32a1-Flp*, which expresses Flp in all GABAergic neurons (**Figure S5A**). We observed robust Ten3^Halo^ signals in the CP (**Figure 5B_3_**), confirming local production of Ten3 in the CP, as also indicated by mRNA *in situ* hybridization analysis (**Figure S6A**). Interestingly, some of the Ten3^Halo^ signals overlapped with Ten3^HA^ signals of the CP (**Figure 5B_4_**). The persistence of Ten3^HA^ could be contributed by (1) expression of Ten3 in non-GABAergic CP neurons, such as cholinergic interneurons^42^; (2) incomplete recombination in GABAergic CP neurons; and/or (3) input axons from cortex and/or thalamus.

**Figure 5.**
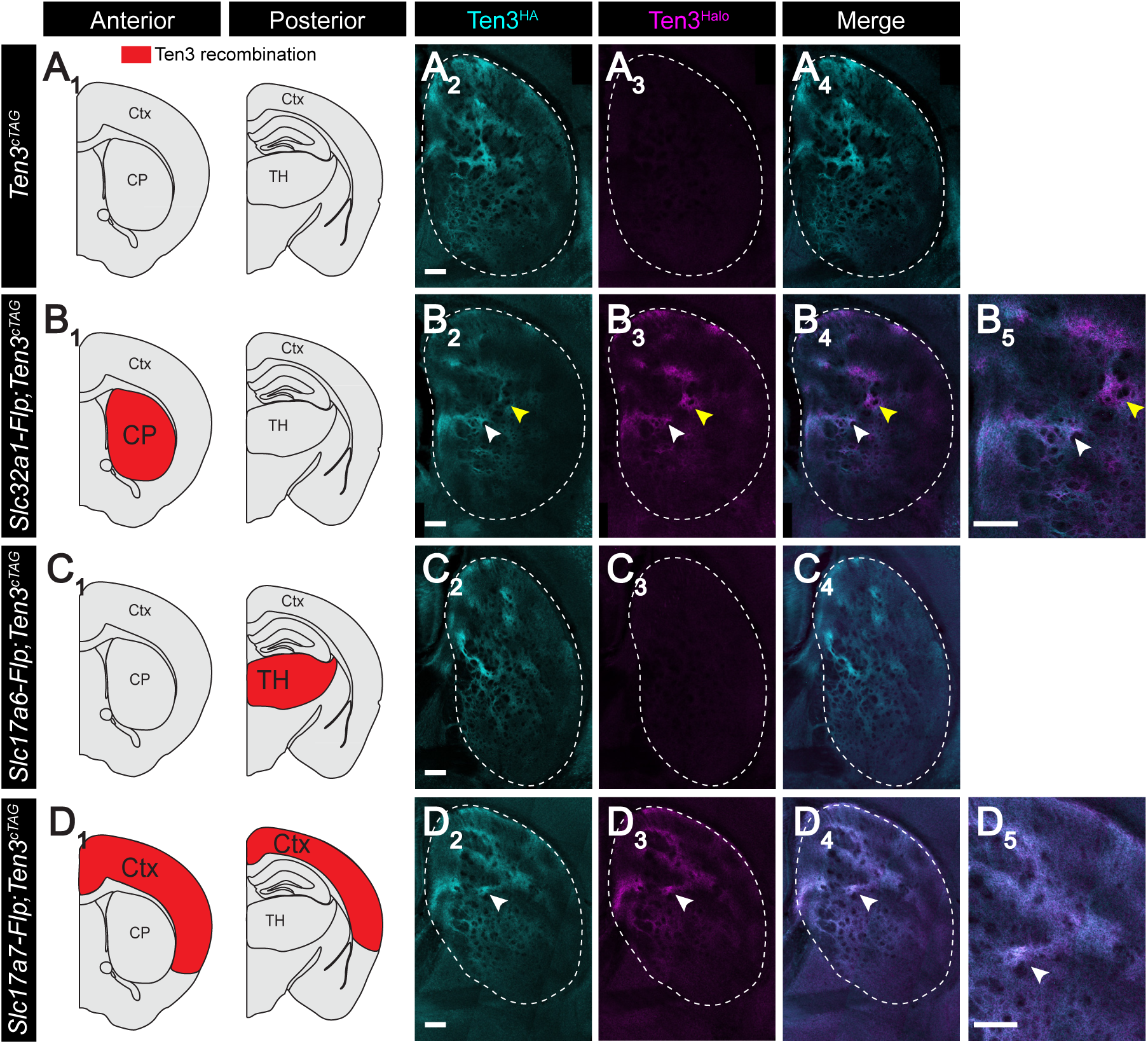
Corticostriatal projections follow the Ten3→Ten3 rule. Left panels are two coronal sections of the mouse brain atlas, with red highlighting regions of recombination due to the expression patterns of the specific Flp driver. Ctx, cortex; TH, thalamus. Right panels are experimental data in P4 mice. CP is demarcated by the dashed outlines. Scale bar, 200 µm. Genotypes on the left. (A) In the absence of Flp driver, *Ten3^cTAG/cTAG^* mice express Ten3^HA^ but not Ten3^Halo^ protein in the CP (dash outlined). (B) In the presence of *Slc32a1-Flp*, which is expressed in all GABAergic neurons, many Ten3^+^ CP regions have strong Halo signal (yellow arrows). Some regions are high for both Halo and HA (white arrows). (C) In the presence of *Slc17a6-Flp*, which is expressed in all excitatory neurons in the thalamus, the lack of Halo signal in CP suggests that thalamic excitatory projection neurons do not express Ten3 in their axon terminals at this stage. (D) In the presence of *Slc17a7-Flp*, which is expressed in all excitatory neurons in the cortex, the overlap between Halo and HA signals in the CP (white arrows) indicates that Ten3^+^ cortical neurons project their axons to Ten3^+^ striatal regions. Scale bar, 200 µm. See Figure S1 and S5 for additional data.

Next, we examined whether thalamic excitatory neurons contribute Ten3 to the CP. To test this, we crossed *Ten3^cTAG^*mice with *Slc17a6-Flp*, which expresses Flp in most/all thalamic excitatory neurons^43,44^ (**Figure S5B**). We did not detect Ten3^Halo^ axon labeling in the CP (**Figure 5C**), suggesting that thalamic axons that innervate the CP do not express Ten3 at their axon terminals at this developmental stage (P4). This contrasts with previous studies suggesting that Ten3 is expressed in both the CP and parafascicular nuclei, where it was proposed to regulate thalamo-striatal connectivity.^45,46^ This discrepancy could reflect differential detecting thresholds for mRNA and protein or post-transcriptional regulation. Although prior studies reported *Ten3* mRNAs in the parafascicular nuclei, we did not detect high levels of Ten3 protein in this region throughout development (**Figure S6G**). While *Ten3* mRNA persist in many brain regions into adulthood^47^, Ten3 protein levels sharply decline after postnatal day 4 in most regions we examined (**Figure S2**), suggesting post-transcriptional regulation.

Finally, to test whether Ten3^+^ cortical excitatory neurons project axons to the CP, we crossed *Ten3^cTAG^*mice with *Slc17a7-Flp*, where Flp is expressed primarily in cortical excitatory neurons but not in any of the CP neurons nor thalamic neurons (**Figure S5C**). Ten3^Halo^ signal was robustly detected in the CP and coincided with Ten3^HA^-expressing regions, including Ten3^HA^ hotspots at the intermediate zone between Ten3^+^ and Lphn2^+^ regions (**Figure 5D**). These data indicate that at least some Ten3^+^ cortical neurons project to the CP, and such projections follow the ‘Ten3→Ten3’ rule. Interestingly, retrosplenial area and posterior parietal association areas of the cortex had high Ten3 expression (**Extended Data 1**) and are known to project to the dorsomedial CP.^48^ These findings suggest that Ten3 expression may contribute to topographic cortical inputs that target distinct CP subregions.

Together, the inverse expression patterns of Ten3 and Lphn2 at CP, both as global gradients and segregated hotspots at the intermediate zone, could be used to organize cortical input and pallidal output of CP neurons following the ‘Ten3→Ten3’ connectivity rule.

### Inverse expression patterns of Ten3 and Lphn2 in both Purkinje cells and cerebellar nuclei reveal new molecular organization of the cerebellum

The cerebellum has a highly organized and stereotyped architecture and plays key roles in motor coordination, sensorimotor integration, and cognition.^49–51^ All information processing in the cerebellar cortex channels through a single layer of Purkinje cells (PCs) projecting their axons to the cerebellar nuclei (CN), which then broadcast to the rest of the brain. The PC→CN projections are established early in development and exhibit precise spatial organization,^49^ yet the molecular mechanisms that guide PC axons to their appropriate nuclear targets remain poorly defined.

In coronal sections of the developing cerebellum, we found that Ten3 and Lphn2 proteins were inversely expressed in PCs in alternating segments across the medial–lateral axis (**Figure 6A**; **Figure S2E**), with sharp transitions (**Figure 6B**). Quantification of normalized fluorescence intensities along the PC layer (**Figure 6A**) revealed the reciprocity of Ten3 and Lphn2 across hemispheres in an individual mouse (**Figure 6C**). Spearman’s correlation analysis confirmed an inverse relationship across different mice (**Figure 6D**). Inverse expression of Ten3 and Lphn2 was also evident in the CN (**Figure 6E**), with preferential expression of Ten3 in dorsomedial and ventrolateral CN, inversely to Lphn2 enrichment in intermediate CN in this coronal section (**Figure 6F**). This pattern was confirmed by Spearman’s correlation analysis across multiple mice (**Figure 6G**). To further assess the stereotypy of Ten3 and Lphn2 expression, we quantified spatial similarity on whole-mount-stained cerebella. This analysis revealed minimal intra-hemispheric overlap between Ten3 and Lphn2, and high inter-hemispheric and inter-animal similarity for Ten3 or for Lphn2 (**Figure S7D**), to a similar degree as other regions with well-established, highly stereotyped expression patterns (**Figure S7E**). Double *in situ* hybridization for *Ten3* and *Lphn2* mRNA revealed similar inverse expression pattern throughout development in both PC layer and CN (**Figure S7A–C**), validating that at least some of the protein expression in both PC and CN is contributed by local neurons.

**Figure 6.**
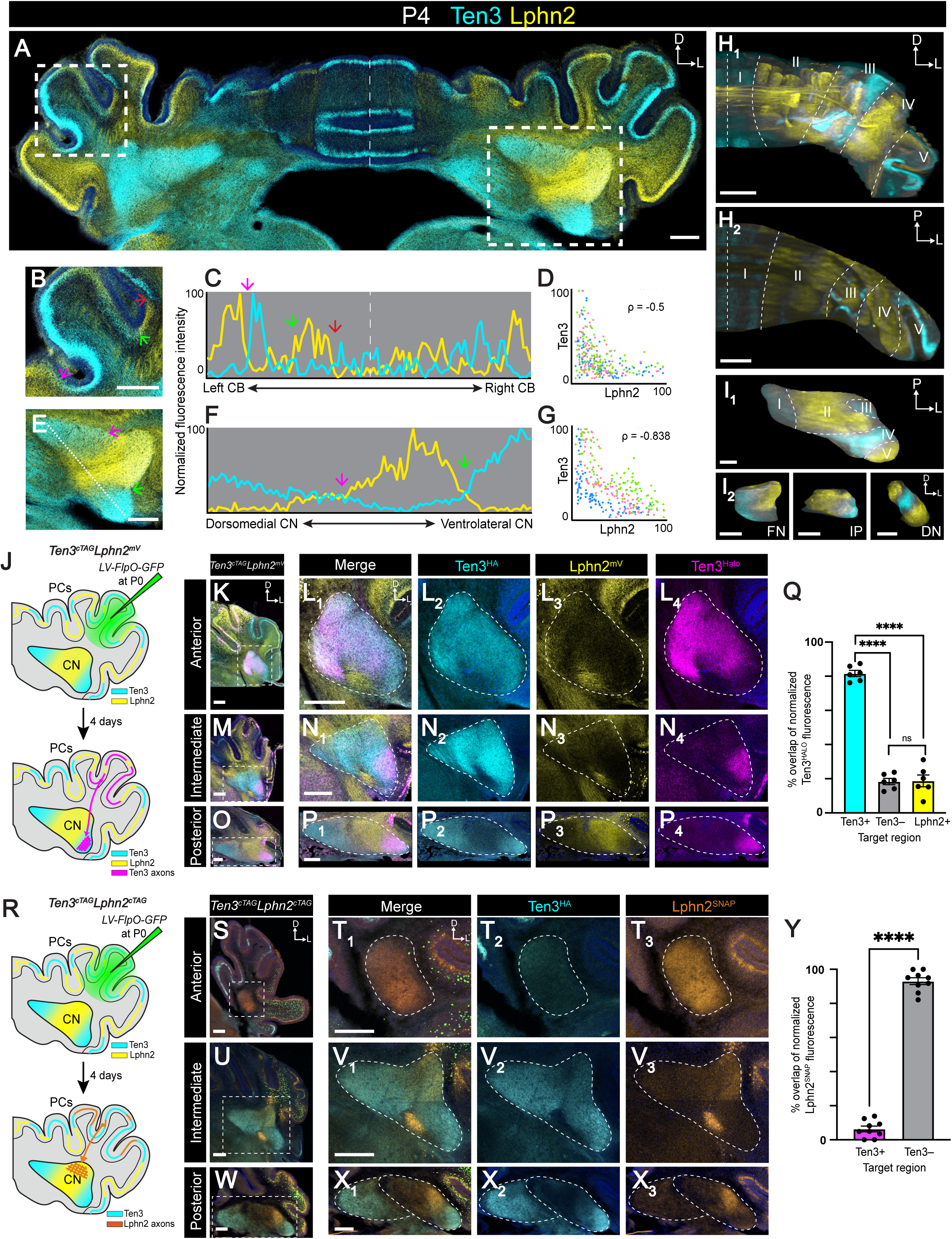
Inverse expression of Ten3 and Lphn2 in both Purkinje cells and cerebellar nuclei. (A) Ten3 and Lphn2 protein expression in the cerebellum of a P4 brain, shown in a coronal section. Dashed vertical line represents the midline. Scale bar, 500 µm. (B) Enlarged view of the cerebellar cortical region outlined in (A, left), with different colored arrows indicating the transitions of Ten3- and Lphn2-expression bands in the Purkinje cell layer. Scale bar, 500 µm. (C) Quantification of normalized Ten3 and Lphn2 protein fluorescence intensity along the PC layer across the left–right axis of the entire cerebellar cortex shown in (A). Different colored arrows indicate the same positions marked by the same colors in (B). (D) Spearman’s correlation between Ten3 and Lphn2 expression in the cerebellar cortex from data in (C); n = 3 mice. Each dot represents the normalized expression level of Ten3 and Lphn2 in a single x-y spatial bin (n = 100 bins per brain along the PC layer). Dots of the same color are from the same mouse (see STAR Methods for detail). Spearman’s correlation, ****p < 0.0001. (E–G) Same as (B–D), except for cerebellar nuclei. Spearman’s correlation, ****p < 0.0001. (H) Ten3 and Lphn2 protein expression of the right hemisphere of the cerebellar cortex at P4 in a 3D volume (H_1_) and a 2D horizontal slab (H_2_), after whole brain clearing and light-sheet imaging. Dashed lines denote each domain. Scale bar, 300 µm. (I) Ten3 and Lphn2 protein expression of the right hemisphere of the cerebellar nuclei at P4 in a 3D volume, after whole brain clearing and light-sheet imaging (I_1_). Ten3 and Lphn2 expression do not align strictly with the fastigial nucleus (FN; I_2_ left), interposed nucleus (IP; I_2_ middle), and dentate nucleus (DN; I_2_ right). Dashed lines denote each domain. Scale bar, 300 µm. (J) Experimental design (top) and summary of results (bottom) for tracing Ten3^+^ PC→CN connectivity. LV, lentivirus. Recombination of *Ten3^cTAG^* allele at the injection site causes Ten3^+^ PCs to switch from expressing Ten3^HA^ to Ten3^Halo^. Recombined Ten3^Halo+^ PC axons target CN regions that express Ten3^HA^. (K, M, O) Low-mag coronal sections of *Ten3^cTAG/cTAG^;Lphn2^mV/mV^* cerebella visualized at P4. Each row indicates an individual mouse. The cerebellar cortices of these mice have been injected at P0 with *LV-FlpO-GFP* (green in K). Ten3^HA^ is in cyan and Ten3^Halo^ in magenta. (L, N, P) Ten3^HA^, Lphn2^mV^, and Ten3^Halo^ protein expression in the anterior (L), intermediate (N), and posterior (P) cerebellar nuclei of *Ten3^cTAG/cTAG^;Lphn2^mV/mV^* mice at P4. Panels L_1-4_, N_1-4_, and P_1-4_ correspond to dashed boxes in panels K, M, O, respectively. Dotted lines outline the CN. (Q) Percent overlap of normalized fluorescence of Ten3^Halo+^ PC axons with Ten3^HA^-positive or -negative, or Lphn2^mV^-positive CN target regions in *Ten3^cTAG/cTAG^;Lphn2^mV/mV^*mice at P4 (n = 6). Paired t-test; ****P < 0.0001; ns, not significant. (R) Experimental design (top) and summary of results (bottom) for tracing Lphn2^+^ PC→CN connectivity. LV, lentivirus. Recombination of *Lphn2^cTAG^* allele at the injection site causes Lphn2^+^ Purkinje cells to switch from expressing Lphn2^FLAG^ to Lphn2^SNAP^. Recombined Lphn2^SNAP+^ PC axons target Ten3^HA^-negative regions of the CN. (S, U, W) Low-mag coronal sections of *Ten3^cTAG/cTAG^;Lphn2^cTAG/cTAG^*cerebella visualized at P4. Each row indicates an individual mouse. The cerebellar cortices of these mice have been injected at P0 with *LV-FlpO-GFP* (green). Ten3^HA^ is in cyan and Lphn2^SNAP^ in orange. (T, V, X) Lphn2^SNAP+^ PC axons target Ten3^HA^-negative region in the anterior (T), intermediate (V), and posterior (X) cerebellar nuclei of *Ten3^cTAG/cTAG^;Lphn2^cTAG/cTAG^* mice at P4. Panels T_1–3_, V_1–3_, and X_1–3_ correspond to dotted box in panels S, U, W, respectively. Dotted lines outline the CN. (Y) Percent overlap of normalized fluorescence of Lphn2^SNAP+^ PC axon projections to Ten3^HA^-positive or Ten3^HA^-negative CN target regions in *Ten3^cTAG/cTAG^;Lphn2^cTAG/cTAG^* mice at P4 (n = 9). Paired t-test; ****P < 0.0001. Scale bar, 200 µm for all panels unless specified. See Figure S1, S2 and S7, and Videos S4 and S5 for additional data.

Previous studies of the cerebellar cortex have extensively characterized PC organization along the medial–lateral axis based on expression patterns of specific genes,^52^ revealing alternating arrays of parasagittal zebrin II-positive (zebrin^+^) and zebrin II-negative (zebrin^−^) stripes that run orthogonal to the medial–lateral axis across the entire cerebellar cortex.^52–54^ Given our finding of Ten3^+^ and Lphn2^+^ ‘bands’ distributed across the medial–lateral axis, we next asked whether these expression patterns correspond to the zebrin stripes. Co-staining of Ten3 and Lphn2 with either zebrin^+^ or zebrin^−^ molecular markers, aldolase C or PlcB4, respectively,^55,56^ we did not find correspondence in their expression patterns (**Figure S7F, G**), indicating that Ten3^+^ and Lphn2^+^ ‘bands’ represent a distinct organization from the zebrin stripes. Indeed, visualizing Ten3^+^ and Lphn2^+^ PCs in 3D revealed that many parasagittal bands of Ten3 and Lphn2 apparent from sections (e.g., **Figure 6A**) were in fact connected in 3D, forming continuous regions (**Figure S7H, Video S4**). The Ten3^+^ and Lphn2^+^ regions in 3D could nevertheless be segmented into five discrete domains per hemisphere (**Figure 6H**). Notably, 3D analysis of expression in the CN also revealed five discrete domains spanning 3D organization of the CN (**Figure 6I_1_, Video S4**). The five CN domains defined by Ten3 and Lphn2 expression do not align strictly with the three histologically separable cerebellar nuclei—namely, the fastigial, interposed, and dentate nucleus (**Figure 6I_2_; Figure S7A)**—suggesting that Ten3 and Lphn2 expression defines a molecular organization distinct from the traditional cytoarchitectural boundaries.

The observed Ten3 and Lphn2 pattern in the PC layer stands in stark contrast to the column-like organization typically associated with parasagittal stripes across all axes (**Video S5**), as most well-known cerebellar molecular markers conform to a stripe-like pattern. The five broad, continuous domains in both PCs and CN defined by inverse Ten3 and Lphn2 expression represent novel molecular heterogeneity in the cerebellum, which may reflect previously unknown functional organization within the cerebellar circuits.

### Purkinje cell→cerebellar nuclei projections follow the ‘Ten3→Ten3, Lphn2→Lphn2’ rule

Given the inverse expression of Ten3 and Lphn2 in both PCs and CN, we next tested whether the PC→CN projection follows the ‘Ten3→Ten3, Lphn2→Lphn2’ rule. We first tested whether Ten3^+^ PC axons overlapped with Ten3^+^ target region of CN. We injected lentivirus expressing *FlpO-GFP* into the cerebellar cortex of *Ten3^cTAG/cTAG^;Lphn2^mV/mV^* mice at P0 and examined Ten3^Halo+^ PC axons in the CN (**Figure 6J**). We found that Ten3^Halo+^ PC axons preferentially projected to the Ten3^HA+^ regions of the CN and avoided Lphn2^mV+^ regions of the CN (**Figure 6K–P**). Quantitative analysis confirmed significantly greater overlap between Ten3^Halo+^ axon signals with Ten3^HA+^ target regions compared to Lphn2^mV+^ target regions (**Figure 6Q**).

To test whether Lphn2^+^ Purkinje cells project to Lphn2^+^ regions of the CN, we utilized our newly generated Lphn2 conditional tag mice (**Figure S1B**) and injected the *FlpO-GFP* lentivirus into the cerebellar cortex of *Ten3^cTAG/cTAG^;Lphn2^cTAG/cTAG^* mice at P0. This would switch Lphn2 expression from Lphn2^FLAG^ to Lphn2^SNAP^ in PC axons, while leaving Ten3^HA^ unrecombined in CN target region (**Figure 6R**). Indeed, Lphn2^SNAP+^ PC axons projected preferentially to Ten3^−^ regions of the CN, avoiding Ten3^HA+^ territories (**Figure 6S–Y**).

In summary, Ten3 and Lphn2 show an inverse expression pattern in both the PCs and CN, and PC→CN projections follow the ‘Ten3→Ten3, Lphn2→Lphn2’ rule. Together with our findings in the visual, auditory, and basal ganglia systems (**Figures 2–5**), as well as the extended hippocampal network,^15,17^ these findings suggest a generalizable mechanism by which spatially segregated Ten3 and Lphn2 could guide the formation of precise long-range connections in parallel networks using the ‘Ten3→Ten3, Lphn2→Lphn2’ connectivity rule.

### Lphn2 is required for targeting precision of PC→CN axons

Given our demonstration that Ten3-mediated homophilic attraction and Lphn2–Ten3–mediated heterophilic repulsion instruct target selection of the extended hippocampal circuit to follow the ‘Ten3→Ten3, Lphn2→Lphn2’ connectivity rule,^15,17^ we asked whether Lphn2 plays a functional role in executing the ‘Ten3→Ten3’ projection rule observed in the cerebellum. We used our conditional tag mice to examine if Ten3^+^ PC axons would mistarget to Ten3^−^ regions of CN when *Lphn2* was deleted.

We injected *FlpO-GFP* lentivirus into PCs of *En1-Cre;Lphn2^+/+^;Ten3^cTAG^*mice (control, **Figure 7A**) or *En1-Cre;Lphn2^fl/fl^;Ten3^cTAG^* mice (experimental, **Figure 7B**) at P0. *En1*, encoding Engrailed homeobox 1, is expressed in developing rhombomere I that gives rise to both PCs and CN. Recombination in both regions was confirmed with a Cre-dependent reporter (**Figure S5D**), which would result in *Lphn2* being deleted in both PCs and CN. Ten3^Halo+^ axons from PCs selectively targeted Ten3^HA^ regions in the anterior, intermediate, and posterior CN in *Lphn2^+/+^*control mice (**Figure 7C–F**). By contrast, Ten3^Halo+^ axons from PCs targeted more broadly into Ten3^−^ CN regions (non-Ten3^HA^ regions) in *Lphn2^cKO^* mice, indicative of mistargeting (**Figure 7G–J**, red arrows). This mistargeting was observed at all levels of the anterior–posterior axis, with variations in severity. Some Ten3^Halo+^ PC axons still innervated Ten3^HA^ domains of the CN, which could be contributed by Ten3–Ten3 homophilic attraction not affected by our experimental manipulation. Additionally, the order in which Ten3^Halo+^ axons encounter Ten3- or Lphn2-expressing domains may influence their targeting as demonstrated in the companion paper^17^, contributing to heterogeneity of the phenotypes.

**Figure 7.**
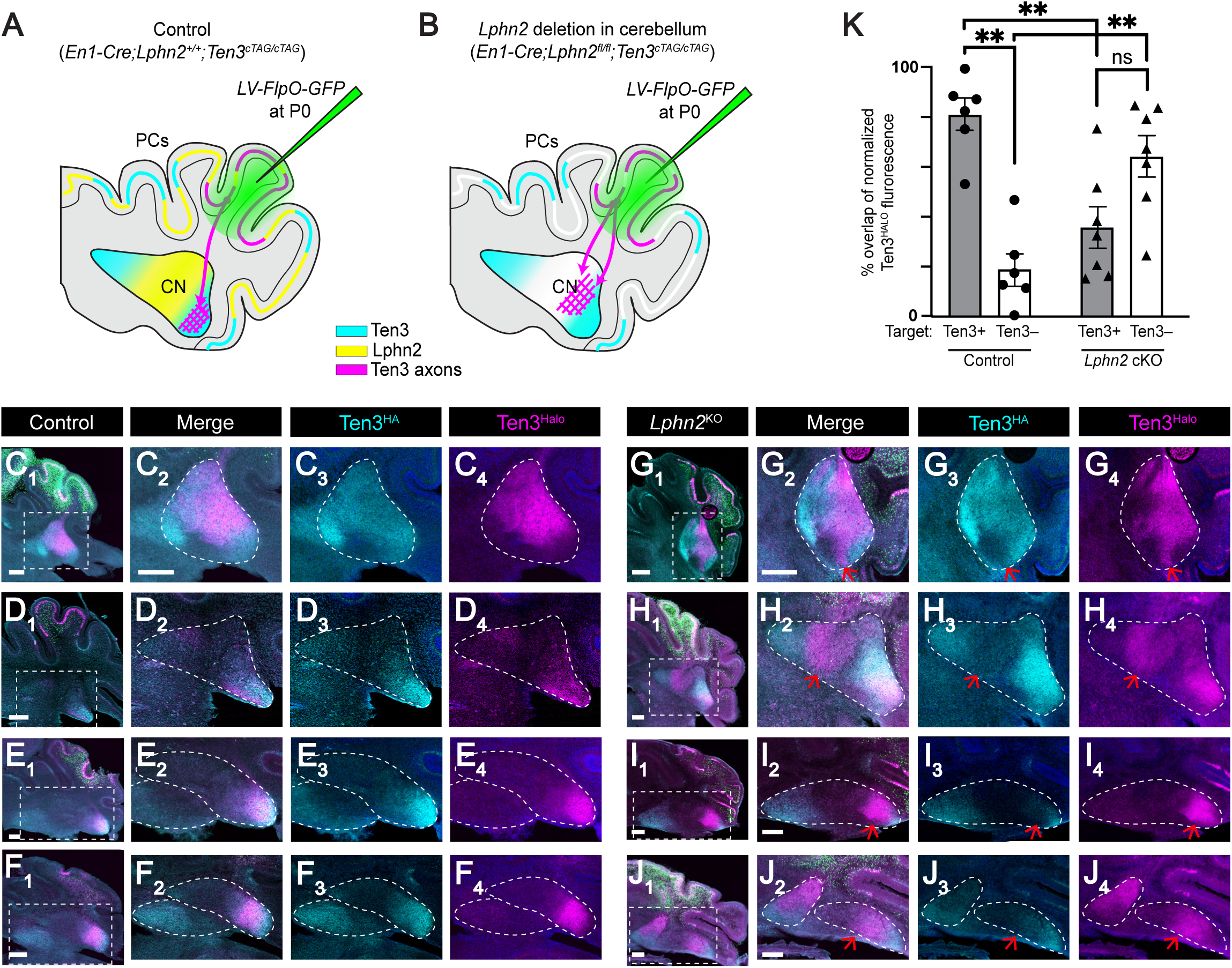
Lphn2 is required for Ten3^+^ Purkinje cell axons to precisely target Ten3^+^ regions of the cerebellar nuclei. (A, B) Experimental design and summary of results for examining Ten3→Ten3 projection in control (A) and *Lphn2* conditional knockout in the cerebellum (B). LV, lentivirus. (C–F) Ten3^HA^ and Ten3^Halo^ protein expression in the anterior (C), intermediate (D), and posterior (E & F) CN of control (*En1-Cre;Lphn2^+/+^;Ten3^cTAG/cTAG^*) mice at P4. (G–J) Ten3^HA^ and Ten3^Halo^ protein expression in the anterior (G), intermediate (H), and posterior (I & J) CN of *Lphn2* conditional knockout (*En1-Cre;Lphn2^fl/fl^;Ten3^cTAG/cTAG^*) mice at P4. Red arrows indicate mistargeting of Ten3^Halo+^ axons in a Ten3^HA^-negative region. (K) Percent overlap of normalized fluorescence of Ten3^Halo+^ Purkinje cell axonal projections to Ten3^HA^-positive or -negative CN target regions in control (n = 6) and *Lphn2* conditional knockout mice (n = 7) at P4. Paired (within the same genotype) and unpaired (between genotypes) t-tests were performed. **p < 0.01. Mean ± SEM. Scale bar, 200 µm for all panels. See Figure S1 and S5 for additional data.

Nevertheless, quantification of axonal overlap overall revealed a significant reduction in Ten3^Halo^ alignment with Ten3^HA^ regions and a significant increase in Ten3^Halo^ overlap in Ten3^−^ CN regions in *Lphn2^fl/fl^*mice compared to *Lphn2^+/+^* control mice (**Figure 7K**). These results indicate that *Lphn2* is required for PC→CN projections to follow the ‘Ten3→Ten3’ rule, likely caused by Lphn2 acting as a repulsive ligand^15,17^ to restrict Ten3^+^ PC axons to appropriate Ten3^+^ CN targets.

## CONCLUDING REMARKS

Here, we describe striking, inverse expression of cell-surface proteins Ten3 and Lphn2 across the visual, auditory, basal ganglia, and cerebellar circuits during development. These inverse expression patterns emerge in three distinct modes: 1) gradients in the retina–superior colliculus of the visual circuit and the cochlear nucleus–superior olivary complex of the auditory circuit (**Figure 2 and 3)**; 2) discrete domains in the cerebellum (**Figure 6**) and the extended hippocampus described in the companion paper;^17^ and 3) a combination of both in the striatal circuit (**Figure 4**). These distinct spatial expression modes may reflect different underlying organizational logics for specific circuits. In retinotopic and tonotopic maps, nearby neurons in the input field project to nearby neurons in the target field to preserve continuous spatial and frequency maps. Continuous gradients of wiring molecules enable their levels to instruct targeting specificity.^1,8,23–26^ By contrast, expression in discrete domains may serve as independent modules for circuit segregation and parallel information processing, as in the case of the medial and lateral subnetworks in the extended hippocampal circuit.^57^ The global gradients of Ten3 and Lphn2 expression in CP may reflect continuous representation of sensorimotor function of the basal ganglia.^37,41^ The discrete hotspots in CP and discrete domains in the cerebellum of Ten3 and Lphn2 expression we discovered do not match with previous described striosomes/matrix or parasagittal zebrin+/zebrin– bands, suggesting novel molecular organizations, the functional significance of which remains to be discovered. These findings also suggest that detailed examination of Ten3 and Lphn2 expression in other circuits (**Extended Data 1**), like what we have done here for the four systems, may reveal previously unknown connection specificity that might lead to new insights into their functional organization.

The temporal dynamics of Ten3 and Lphn2 expression suggest a widespread role in target selection. In systems where timing of target selection of axons has been determined, this timing overlaps with peak expression of Ten3 and Lphn2 (**Figure S2**). This spatiotemporal regulation of CSP expression may allow a limited set of molecules to direct wiring of many circuits across diverse brain regions.

Interestingly, the inverse expression of Ten3 and Lphn2 is observed across a diverse array of brain regions, including the forebrain, midbrain, and hindbrain, despite their distinct developmental origins. This raises an intriguing question about whether a conserved regulatory mechanism governs this expression pattern across the brain, or if region-specific cues independently establish similar molecular expression patterns. Regardless of details of the regulatory mechanisms, our findings suggest that tweaking the expression patterns of key existing wiring molecules is an effective strategy for accommodating the increasing complexity of the nervous system during evolution.

Together with our findings in the companion manuscripts demonstrating functional roles of Ten3 and Lphn2 in instructing target selection in the extended hippocampal circuit^17^ and somatotopic map^21^, these results define a conserved molecular logic for target recognition that is used repeatedly across the nervous system to establish precise neural circuits. The inverse expression of Ten3 and Lphn2, along with their molecular function as a ligand–receptor pair mediating both attraction and repulsion, suggests a global molecular strategy by which distinct circuits achieve precise circuit assembly throughout development.

## RESOURCE AVAILABILITY

### Lead contact

Requests for further information and resources should be directed to and will be fulfilled by the lead contact, Liqun Luo.

### Materials availability

Mouse lines generated in this study will be deposited to the Jackson Laboratory.

### Data and code availability

- Interactive platform (Extended Data 1. Interactive visualization of Ten3 and Lphn2 expression in horizonal sections of E17 and P2 mouse brains, related to Figure 1) will be deposited at the Stanford Digital Repository and will be publicly available as the date of publication at https://doi.org/10.25740/jp473xf0650. To visualize the datasets, download all files. To visualize E17 data, open the file *Ten3_Lphn2_E17.tif* in Fiji by dragging it into the Fiji console. Next, launch the ROI manager in Fiji (*Analyze* > *Tools* > *ROI Manager*) and load the file *E17_ROIs.zip*. Each ROI corresponds to a defined region of interest: clicking on the ROI names will allow you to view Ten3 and Lphn2 expression across respective regions. Same for P2. Ten3 is in cyan and Lphn2 is in yellow.
- This paper does not report original code.
- Any additional information required to reanalyze the data reported in this paper is available from the lead contact upon request.

## ACKNOWLEDGEMENTS

We thank members of the Luo Lab, especially Ellen Gingrich, Tom Hindmarsh-Sten, Lijun Qi, Zhuoran Li, Alex Starr, Yunming Wu, and Cheng Lyu, as well as Julia Kaltschmidt, Jennifer Raymond, and Kang Shen for their support, expertise, and insights on this study. We also thank Caiying Guo and the Transgenic Facility at Janelia Research Campus for generating the cTAG mice; Yiming Zhang, Huyan Meng, and Zhigang He at the Boston Children’s Hospital Viral Core for the *LV-FlpO-GFP* virus; Artur Kania and Kevin Sangster for feedback on the manuscript; Kenneth Magpusao and Carlota Manalac for animal assistance; David Luginbuhl and Mary Molacavage for administrative assistance. D.T.P. was supported by the American Australian Association Education Fund Scholarship. L.L. is an HHMI investigator. This work was supported by the National Institutes of Health (R01-NS050580 to L.L.), and the Simons Foundation (SFARI Fellows-to-Faculty Award to D.T.P.)

## AUTHOR CONTRIBUTIONS

L.L., D.T.P., and U.C. conceived the project and designed all experiments. U.C. and D.T.P. performed majority of the experiments and data analyses. J.H.S. performed whole brain registration and Dice coefficient analyses, and Y.Z. conducted the *in situ* hybridization assays. I.R. assisted with the iDISCO experiments. L.L. and D.T.P. supervised the study. U.C., D.T.P., and L.L. wrote the manuscript.

## DECLARATION OF INTERESTS

The authors declare no competing interests.

### STAR Methods

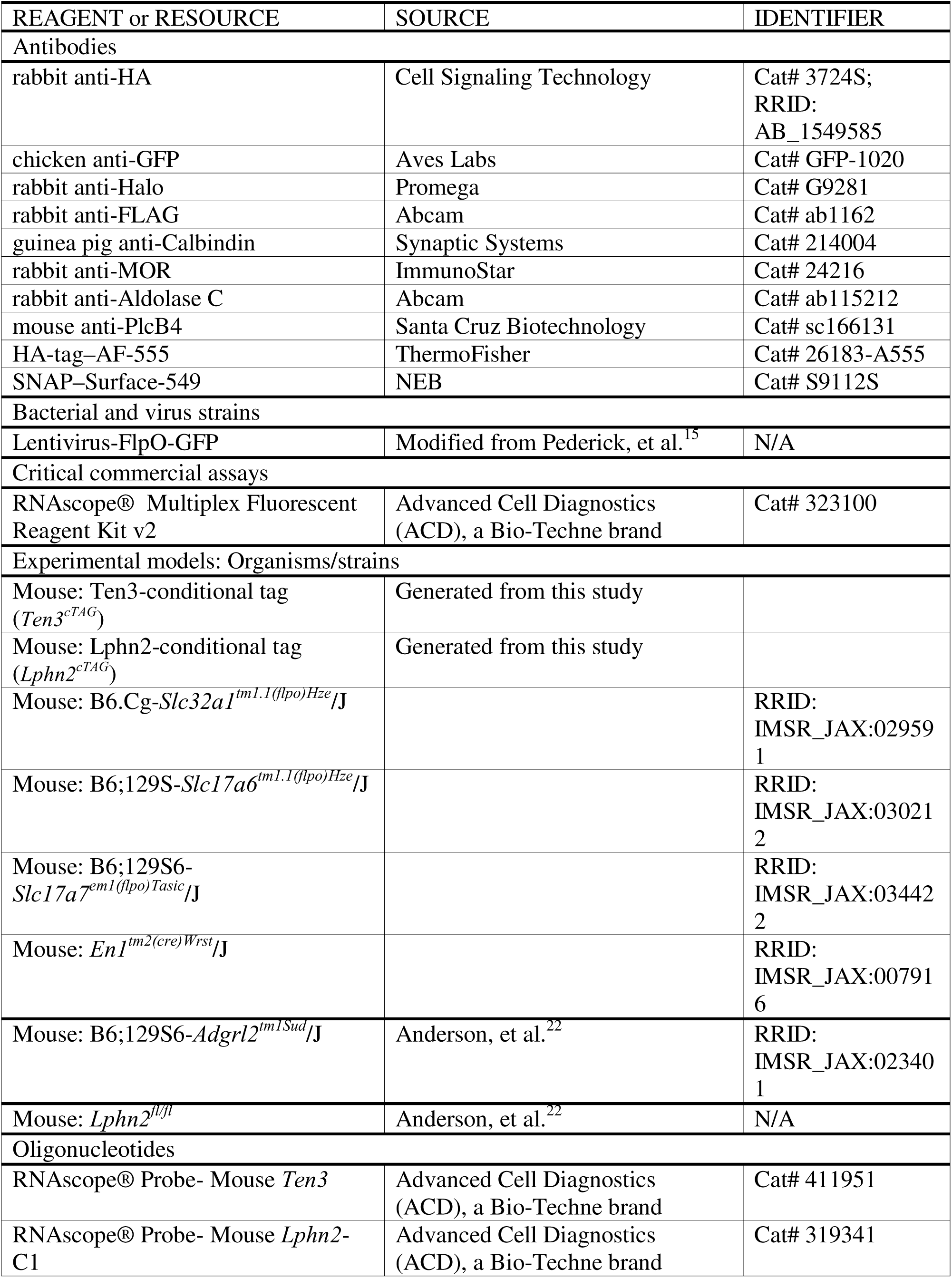
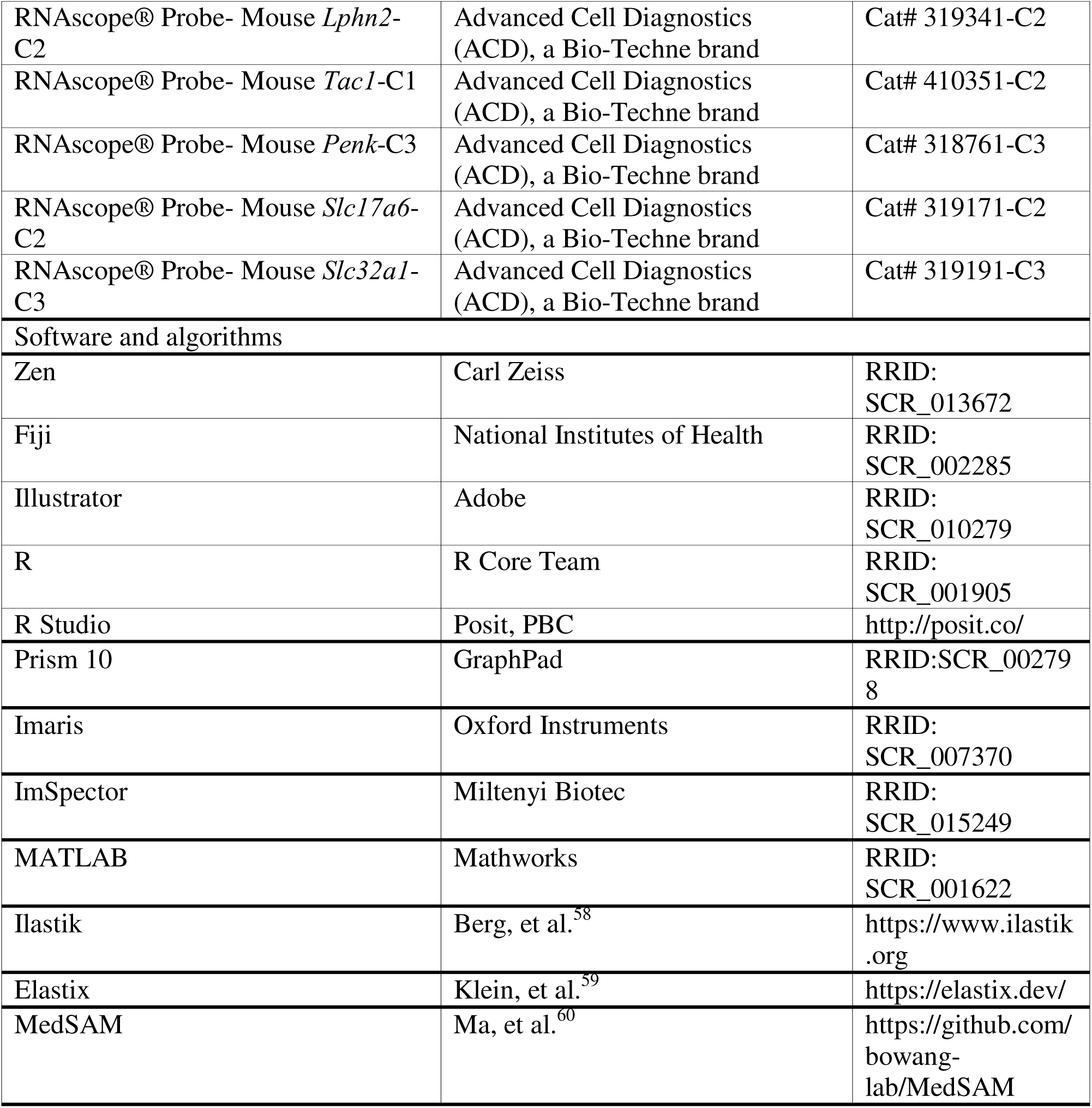
Key resources table.

## EXPERIMENTAL MODEL AND PARTICIPANT DETAILS

All animal procedures followed animal care guidelines approved by Stanford University’s Administrative Panel on Laboratory Animal Care and Administrative Panel on Biosafety in accordance with NIH guidelines. Both male and female mice were used, where they were group housed with access to food and water *ad libitum*. Mice at embryonic day (E) 15 and 17, and postnatal day (P) 0, 2, 4, 8, 21, 42 were used.

*Slc32a1-Flp* (Jackson Laboratory, Stock 029591), *Slc17a6-Flp* (Jackson Laboratory, Stock 030212), and *Slc17a7-Flp* (Jackson Laboratory, Stock 034422) were bred to used *Ten3^cTAG^* to switch Ten3 tags in specific cell types. *En1-Cre* (Jackson Laboratory, Stock 007916) mice^61^ as well as *Lphn2^mVenus^* and *Lphn2^fl/fl^* mice^22^ were on a C57BL/6 and 129 mixed background and were backcrossed to wild-type CD1 mice to improve pup survival rate. *Ten3^cTAG^* and *Lphn3^cTAG^* mice were generated in this study. All genotyping was performed as previously described^13,15,22,61^.

## METHOD DETAILS

### Generation of conditional tagged mice

*Ten3* and *Lphn2* conditional tag (cTAG) mice were generated by the Gene Targeting and Transgenics core at HHMI/Janelia Research Campus. Both mouse lines were generated by homologous recombination in ES cells using standard procedures. The *Ten3^cTAG^* construct was designed such that exon 29 was fused in-frame with an FRT-HA tag, followed by an SV40 polyA signal and an FRT-HaloTag. A self-excising selection cassette containing a loxP2272-flanked RNA polymerase II promoter-NeoR and ACE-Cre was inserted downstream of the Halo tag. This cassette is removed automatically in F1 mice. Homology arms of 4298 bp (5’) and 2231 bp (3’) were retrieved from BAC clone RP23-369L1 using recombineering techniques. To enhance targeting efficiency, the CRISPR-Cas9 system was employed. The construct was co-electroporated into an F1 hybrid 129S6 × C57BL/6J ES cell line along with a guide RNA (gRNA) 5’-ATCGGCAAGAGGTAACCCCC-3’ and Cas9 protein (Fisher Scientific, Cat# A36498). Forty-eight G418-resistant ES colonies were isolated and screened via nested PCR using primers located outside and inside the construct. The primers used for ES cell screening were as follows: **5’ Arm:** Forward primers: *Ten3 Scr F1*: 5’-ATGGCTTTCGGAGTCACATGC-3; *Ten3 Scr F2*: 5’-TCTTAGTCTCCACCTGAGCA-3’ and Reverse primers: *SV40 PA R*: 5’-GTGGTTTGTCCAAACTCATCA-3’; *Ten3 Scr 5R1*: 5’-CATCGTATGGGTAGAAGTTCC-3’; **3’ Arm:** Forward primers: *ACE Scr R1*: 5’-ACAGCACCATTGTCCACTTG-3’; *ACE Scr R2*: 5’-GCTGGTAAGGGATATTTGCC-3’ and Reverse primers: *Ten3 Scr R1*: 5’-AGGAGACTCTCCAGATCTGT-3’; *Ten3 Scr R2*: 5’-CTGTCATGTGGTACACGCTA-3.’

Three of the 39 ES clones that tested positive for both homology arms were used to generate mice. Chimeric mice were produced by aggregating these ES cells with 8-cell-stage CD-1 embryos. Correct targeting was further confirmed by homozygosity testing in the progeny. All three ES clones contributed to the germline and produced homozygous pups. Chimeras were bred with wild-type C57BL/6J females.

The *Lphn2^cTAG^* construct was designed such that exon 24 was fused in-frame with an FRT-Flag tag, followed by an SV40 polyA signal and an FRT-SNAP tag. A self-excising selection cassette containing a loxP2272-flanked RNA polymerase II promoter-NeoR and ACE-Cre was inserted downstream of the SNAP tag. This cassette is removed automatically in F1 mice. Homology arms of 3686 bp (5’) and 2814 bp (3’) were retrieved from BAC clone RP23-283E24 using recombineering techniques. To enhance targeting efficiency, a CRISPR-Cas9 system was employed. The construct was co-electroporated into an F1 hybrid 129S6 × C57BL/6J ES cell line along with a guide RNA (gRNA) 5’-AGTCTTTAATAGTACGGCTAAGG-3’ and Cas9 protein (Fisher Scientific, Cat# A36498). Forty-eight G418-resistant ES colonies were isolated and screened via nested PCR using primers located outside and inside the construct. The primers used for ES cell screening were as follows: **5’ Arm:** Forward primers: *Lphn2 Scr F1*: 5’-AGGACTGACATCACATGGTC-3’; *Lphn2 Scr F2*: 5’-GCTTCTGGAATGAGCTAGCT-3’ and Reverse primers: *SV40 PA R*: 5’-GTGGTTTGTCCAAACTCATCA-3’; *Flag tag R*: 5’-CGTCGTCATCCTTGTAATCG-3’; **3’ Arm:** Forward primers: *ACE Scr R1*: 5’-ACAGCACCATTGTCCACTTG-3’; *ACE Scr R2*: 5’-GCTGGTAAGGGATATTTGCC-3’ and Reverse primers: *Lphn Scr R1*: 5’-ACACTTAAGAGTGCCTGTGC-3’; *Lphn Scr R2*: 5’-TTCACAGACATGTTTGCACC-3.’

Three of the 25 ES clones that tested positive for both homology arms were used to generate mice. Chimeric mice were produced by aggregating these ES cells with 8-cell-stage CD-1 embryos. Correct targeting was further confirmed by homozygosity testing in the progeny. All three ES clones contributed to the germline and produced homozygous pups. Chimeras were bred with wild-type C57BL/6J females.

### Genotyping

Primers used for genotyping *Ten3^HA^* and *Ten3^Halo^* alleles include Ten3-HA-F (*CTACGATGGGTACTACGTAC*), Ten3-HA-R (*GTGGTTTGTCCAAACTCATCA*), Ten3-Halo-F (*GGTCTGAATCTGCTGCAAGA*), and Ten3-R (*TCCAGTCCAATTGAGTGCAG*). The following oligonucleotide combinations were used to detect the presence of different alleles: Ten3-HA-F/Ten3-HA-R/Ten3-R (*WT*: 329bp; *Ten3^HA^*: 218bp), and Ten3-Halo-F/Ten3-R (*Ten3^Halo^*: 361bp). Primers used for genotyping *Lphn2^FLAG^* and *Lphn2^SNAP^* alleles include Lphn2-FLAG-F (*AAGTATGCCCAACCTTGGAG*), Lphn2-FLAG-R (*CGTCGTCATCCTTGTAATCG*), Lphn2-SNAP-F (*GGCTGGGTTAATGTCGACAT*), and Lphn2-R (*TCACAGGACTGTGAGGTTCA*). The following oligonucleotide combinations were used to detect the presence of different alleles: Lphn2-FLAG-F/Lphn2-FLAG-R/Lphn2-R (WT: 324bp; *Lphn2^FLAG^*: 221bp), and Lphn2-SNAP-F/Lphn2-R (*Lphn2^SNAP^*: 223bp).

### Whole brain immunolabeling and clearing

Mice were injected with 2.5% avertin and were transcardially perfused with cold phosphate buffered saline (PBS) and 4% paraformaldehyde (PFA). Dissected brains were post-fixed with 4% PFA for at room temperature for 1 hour. Whole brain immunolabeling and clearing was performed using a modified iDISCO protocol.^62,63^ Brains were dehydrated through a graded methanol (MeOH) series in B1N buffer, at 20%, 40%, 60%, and 80% MeOH, followed by three washes in 100% MeOH. The incubation time for the dehydration steps were 45 minutes for embryonic brains and 1 hour for postnatal brains. Brains were then incubated overnight in a 2:1 dichloromethane (DCM):MeOH mixture at room temperature. On the next day, they were washed in 100% DCM and 100% MeOH (three times), before incubation in 1:5 H_2_O_2_/MeOH mixture at room temperature. The brains then went through rehydration in the reverse MeOH gradient in B1N buffer. The brains were then incubated in 5% DMSO/0.3M Glycine/PTxwH (10% Triton X–100, Tween-20, heparin, 5% NaN_3_ in PBS) twice for efficient penetration of primary antibodies.

For immunostaining, brains were incubated with primary antibodies—rabbit anti-HA (1:300, Cell Signaling Technology, 3724S) and chicken anti-GFP (1:1000, Aves Labs, GFP-1020) in PTxwH for 5 days (embryonic brains, E15 and E17), 8 days (early postnatal brains, P0–P4), or 11 days (late postnatal and adult brains, P8–P42) brains, at 37°C. After three days of washing in PTxwH, the brains were then transferred into secondary antibodies– donkey anti-rabbit Alexa Fluor 555 (ThermoFisher, A32794) and donkey anti-chicken Alexa Fluor 647 (ThermoFisher, A78952) at 37°C for 4, 6, or 8 days for embryonic, early postnatal, or late postnatal/adult brains, respectively. Following secondary antibody incubation, the brains were washed in PTxwH for 3 days and PBS for 1 day.

For tissue clearing, brains were dehydrated through a graded MeOH series in water, washed three times in 100% MeOH, and incubated overnight in 2:1 DCM:MeOH mixture. Samples were then washed three times in 100% DCM before being cleared in dibenzyl ether (DBE). The cleared brains were subsequently washed daily in ethyl cinnamate for a minimum of three days prior to imaging.

### Light–sheet imaging, registration, and dice coefficient analysis

Cleared whole brains were imaged using UltraMicroscope Blaze light–sheet microscope (Miltenyi Biotec) at 1× magnification. We acquired a series of images using three channels (488 nm for autofluorescence, 560 nm for Ten3, and 640 nm for Lphn2) with dynamic focusing for Ten3 and Lphn2 channels at a z-step size of 3 μm. Autofluorescence channel was imaged without dynamic focusing. Voxel sizes were 3.55 μm × 3.55 μm for E15 brains, and 5.91 μm × 5.91 μm for E17 and postnatal brains. Raw image stacks were first downsampled four times, resulting in a voxel size of 23.64 μm × 23.64 μm × 12 μm. Local Ten3- and Lphn2-positive voxels were binarized using Ilastik.^58^ We note that the segmentation models were optimized for each region of interest (ROI) by training them with local signal information.

For registration, we first chose one brain from the seven imaged brains as our reference brain. This brain had the qualitatively cleanest signals (e.g., least artifacts and deformation). On the right hemisphere of the reference brain, 3-dimensional ROI masks were prepared using a transformer-based segmentation tool^60^ with a combination of the three imaging channels. The other six brains were then registered to this reference brain via non-linear transformation.^59^ For inter-hemisphere analysis within the same brains, we reflected the brain with respect to its anterior–posterior axis and performed registration against the non-reflected brain to minimize variations in region boundaries. Finally, signal overlaps were quantified using Dice coefficients (MATLAB).

### Immunohistochemistry

Mice were injected with 2.5% Avertin and were trascardially perfused with PBS followed by 4% PFA. Dissected brains were post-fixed with 4% PFA at room temperature for 1 hour, and then cryoprotected in 30% sucrose for 2-3 days. Brains were then embedded in OCT (Tissue-Tek), frozen in isopentane bath that was cooled on dry ice and stored at –20°C until sectioning. Using a cryostat, the frozen brains were sectioned at 60 µm thickness into PBS and stored at 4°C. Immunohistochemistry were performed as previously described.^15^ Briefly, the sections were incubated in 10% NDST (0.3% Triton X–100 in PBS and 10% normal donkey serum) for two hours at room temperature, followed by incubation with primary antibodies in 5% NDST (0.3% Triton X–100 in PBS and 5% normal donkey serum) for two nights at 4°C. After multiple washes in PBST (0.3% Triton X–100 in PBS), the sections were incubated with secondary antibodies (1:500) in 5% NDST and DAPI. Sections were mounted with Fluoromount-G (SouthernBiotech, 0100-01) after multiple washes in PBST and PBS. Primary antibodies used were rabbit anti-HA (1:300, Cell Signaling Technology, 3724S), chicken anti-GFP (1:1000, Aves Labs, GFP-1020), rabbit anti-HaloTag (1:500, Promega, G9281), rabbit anti-FLAG-tag (1:1000, Abcam, ab1162), guinea pig anti-Calbindin (1:500, Synaptic Systems, 214004), rabbit anti-MOR (1:500, ImmunoStar, 24216), rabbit anti-Aldolase C (1:300, Abcam, ab115212), and mouse anti-PlcB4 (1:500, Santa Cruz Biotechnology, sc166131). Monoclonal antibody HA-tag–AF-555 (1:300, ThermoFisher, 26183-A555) and SNAP–Surface-549 (1:300, NEB, S9112S) were used. For rabbit anti-FLAG-tag, the antibody was pre–adsorbed on wild–type tissue prior to performing standard immunohistochemistry. Pre-adsorption followed the same immunohistochemistry procedure as stated above. Secondary antibodies conjugated to Alexa 488, 567, Cy3, or 647 (Jackson ImmunoResearch) were used at 1:500 from 50% glycerol stocks. Sections were imaged using the tile scan function on Zeiss LSM 900 confocal microscope with 10× magnification.

### Double *in situ* hybridization

Pups of postnatal day 4 (P4) or younger were anesthetized on ice for 1–2 minutes, whereas older mice were injected with 2.5% Avertin. Mice were euthanized by decapitation and were rapidly dissected into cold PBS. The brains were then embedded into Optimum Cutting Temperature (OCT, Tissue–Tek), snap frozen in dry ice–cooled isopentane and stored in –20°C until sectioning. All samples were processed within 1 minute of dissection to preserve RNA integrity. The fresh–frozen brains were sectioned at 10 µm thickness using a cryostat and mounted onto Superfrost Plus slides (VWR). In situ hybridization was performed using the RNAscope® Multiplex Fluorescent Reagent Kit v2 (ACD Bio) following the manufacturer’s protocol.^64^ Briefly, sections were fixed in 4% PFA for 2 hours at RT, dehydrated in graded ethanol, and treated with Protease IV. Following target-specific probes from ACD Bio were used: *Ten3* (411951), *Lphn2*-C1 (319341), *Lphn2*-C2 (319341-C2), *Tac1* (410351-C2), *Penk* (318761-C3), *Slc17a6* (319171-C2), and *Slc32a1*(319191-C3). The sections were hybridized with these probes for 2 h at 40°C, followed by signal amplification and detection using Opal fluorophores (Akoya Biosciences; Opal 520, 570, and 650; 1:1500 dilution). Sections were counterstained with DAPI, washed, and mounted in ProLong Gold Antifade Mountant (ThermoFisher, P36930). Sections were imaged using the tile scan function on Zeiss LSM 900 confocal microscope (10× magnification) with identical acquisition settings across samples in the same batch of experiments.

### Lentivirus production

Lentivirus expressing Cre-GFP under the ubiquitin-C (UBC) promoter that was originally generated by the Neuroscience Gene Vector and Virus core at Stanford University^15^ was modified such that Cre-GFP was removed and FlpO-Green Lantern was inserted immediately downstream of the UBC promoter using Gibson assembly. Correct insertion of plasmid was verified by whole plasmid sequencing (Plasmidsaurus).

Recombinant vectors were generated by transient transfection of 293T cells. The supernatant was harvested 72 hours post-transfection, filtered through a 0.45 µm membrane, and concentrated via ultracentrifugation (18,000 rpm, 2 h, 4°C; SW 32 Ti rotor). Pellets were pooled, resuspended in 3.5 mL DPBS, and further purified by sucrose cushion ultracentrifugation (20% sucrose cushion, 18,000 rpm, 2 h, 4°C; SW 55 Ti rotor). Final pellets were resuspended in DPBS containing 0.001% pluronic F-68, aliquoted (10 µL), and stored at −80°C.

### Stereotactic injections in neonatal mice

All stereotactic injections were performed on P0 mice. Pups were anesthetized by hypothermia on ice for 2–3 minutes, placed in the stereotaxic frame (Stoelting), and injected with 500 nL (striatum) or 300 uL (cerebellum) of virus per site at 100 nL/min using a pulled glass micropipette (Drummond). *LV-FlpO-GFP* was diluted 1:10 from the original viral stock (1.01 × 10^13^ copies per mL, Boston Children’s Hospital Vector Core). Injection coordinates were determined relative to lambda. Striatum injections were 2.0 mm anterior, 1.3 mm lateral, and 1mm ventral from lambda. Cerebellum injections were 3 mm and 2.8 mm anterior (2 injections), 1.8 mm lateral, and 0.5 mm ventral from lambda. After injection, pups were placed on a warm heating pad for recovery before being returned to their home cage. All injections were verified by *post hoc* analysis.

## QUANTIFICATION AND STATISTICAL ANALYSIS

### Image Analysis

For spatial quantification of Ten3 and Lphn2 in SC (Figure 2G) and the auditory brainstem nuclei (Figures 3C, E), a 200-pixel-wide segmented line was drawn through the cell body layer from each end of the region of interest. These areas were defined by anatomical features or DAPI staining. Segmented lines were drawn, straightened using the Straighten function, background subtraction was performed using the Subtract function and intensity values were measured using the Plot Profile command (performed in Fiji). The intensity plots were resampled into 100 equal bins (MATLAB) and each individual trace was normalized to a maximum of 100. Traces from each mouse were averaged, normalized to a maximum of 100 again and then the normalized average trace from each mouse was combined to give the final distribution.

Due to the unique anatomical constraints and expression characteristics of CP, GPe, and the cerebellum, different quantification methods were used. Fluorescence intensity profiles for Ten3 and Lphn2 were obtained from coronal sections of P4 brains in the CP and GPe (Figure 4B, C), as well as the Purkinje cell layer and the cerebellar nuclei (Figure 6B, E). For each image, ROIs were manually delineated based on anatomical boundaries visible in the DAPI channel. Axes used for quantification were indicated using the dotted lines in Figures 4B, 4C, and 6E. The Purkinje cell layer across the left–right axis of the entire cerebellar cortex was quantified. Mean fluorescence intensity values were extracted along these axes for both Ten3 and Lphn2 channels. To enable direct quantitative comparison across animals, intensity profiles were resampled into 100 equally spaced bins along the axis using a custom MATLAB code previously reported (MathWorks) ^11^. Each bin represented the normalized mean fluorescence intensity within its corresponding spatial segment. These binned intensity values were then used for correlation analysis. Spearman’s correlation coefficients were computed in R, with each brain region analyzed independently (n = 3 mice each). Data points in the scatterplots correspond to a single x–y bin, and individual mice are indicated by distinct colors.

For relative expression analysis in Figure S2 the following method was used. Sections were stained in three batches, with each batch containing one animal of each age. All images from a region of interest in a batch were imaged using a Zeiss LSM 900 confocal microscope on the same day using the same settings. To quantify relative expression levels regions of interest were drawn using anatomical features, and fluorescent intensity values were obtained using the Histogram tool (performed in Fiji). The mean expression level was calculated for the top 10% of pixels in each image and averaged with all other values from the same region of that animal. Averages were then normalized to the age with the highest expression value. Normalized values were grouped by age and region and plotted in Prism 10 (GraphPad).

### Ten3→Ten3, Lphn2→Lphn2 projection mapping analysis

Mice were only included if they met the following criteria: 1) lentivirus injections must be specifically in the striatum for experiments described in Figure 5, or the cerebellar cortex for experiments described in Figure 7; 2) axons projecting to the target region must be at the border of target Ten3–Lphn2 expression; 3) more than three sections per animal needs to have sufficient expression of Ten3^Halo^ or Lphn2^SNAP^ axons at the target region for quantification. Mice that fulfilled these criteria are reported in Figures 5 and 7 and were included in quantifications. Confocal images of sections were acquired using tile function on the Zeiss LSM 900 confocal microscope with identical acquisition settings for all samples within an animal. To account for variability of injection sites between mice, exposure settings were adjusted individually to prevent signal saturation. Projection patterns were quantified using ImageJ/Fiji.

To quantify the overlap of Ten3^Halo+^ or Lphn2^SNAP+^ axons at Ten3^HA+^ target region, ROIs were manually delineated on the DAPI channel using anatomical landmarks. These ROIs were extracted using the clear outside function in ImageJ/Fiji. For each channel, false-positive values were subtracted, and signal thresholds were individually adjusted. A Gaussian blur (radius = 10) was applied, after which ROIs were converted to masks.

Image calculator was then used to compute the number of overlapping pixels across all signal channels. Statistical analyses were performed using paired t-tests using Prism 10 (GraphPad).

### Lphn2-mediated PC→CN wiring analysis

Overlap of Ten3^Halo+^ axons and Ten3^HA+^ target region in *En1-Cre;Lphn2^+/+^;Ten3^cTAG/cTAG^* and *En1-Cre;Lphn2^fl/fl^;Ten3^cTAG/cTAG^*mice were quantified using the same inclusion criteria, imaging procedures, and imaging processing methods described above. Following mask conversion, the Halo channel region was scaled down by 75% of its original size and shape from the center of the mask, and the outer 25% area along the Ten3–Lphn2 border was isolated and extracted using the clear function. An equivalent area—matching the pixel width of the outer 25% region—was also taken from the other side of the Ten3–Lphn2 border. Quantifications of axon innervation were performed using the areas on both sides of the Ten3–Lphn2 border. Corresponding areas in other channels were extracted, and Image Calculator was used to determine the number of overlapping pixels across all signal channels. Statistical analyses were performed as previously described, and all quantification was performed by an experimenter blinded to genotype.

## SUPPLEMENTARY MATERIALS

**Video S1. Three-dimensional reconstruction of whole brain Ten3 and Lphn2 expression at postnatal day 2 (P2), related to Figure 1**.

**Video S2. Horizontal section series showing whole brain Ten3 and Lphn2 expression at embryonic day 15 (E15), E17, postnatal day 0 (P0), P2, P4, P8, related to Figure 1. Anterior is to the left. Each flythrough runs from dorsal to ventral.**

**Video S3. Three-dimensional reconstruction of Ten3 and Lphn2 expression in the caudate putamen (CP) and external segment of the globus pallidus (GPe) at P4, related to Figure 4**.

**Video S4. Three-dimensional reconstruction of Ten3 and Lphn2 expression in the cerebellar cortex and cerebellar nuclei at P4, related to Figure 6**.

**Video S5. Three-dimensional reconstruction of zebrin expression in the adult cerebellar cortex, related to Figure 6**.

Videos S1–4 display Ten3 in cyan and Lphn2 in yellow; for Video S5, zebrin is shown in green.

