## Supplemental Figures for "Inverse expression of Ten3 and Lphn2 across the developing mouse brain suggests a global strategy for circuit assembly"

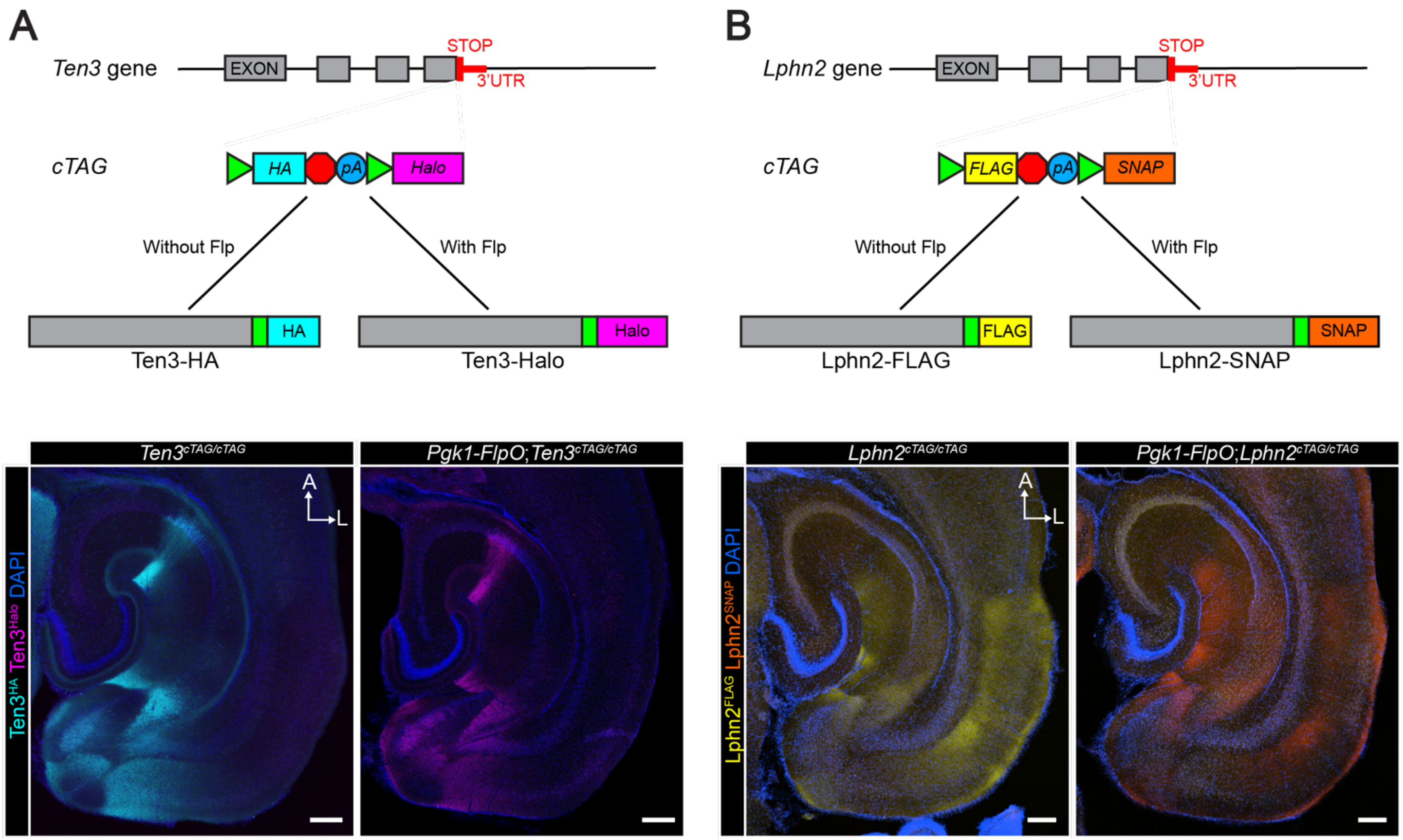


**Figure S1. Design of *Ten3*$^{cTAG}$ and *Lphn2*$^{cTAG}$ mice, related to Figures 1, 3, 5–7**

(A) Schematic of the *Ten3*$^{cTAG}$ mice. To generate the *Ten3* conditional tag mice, an HA epitope tag was added immediately upstream of the STOP codon (red octagon) of the *Ten3* open reading frame, flanked by two *FRT* sites (green triangle), followed by a Halo tag. In the absence of Flp recombinase, Ten3 protein is tagged with HA; in the presence of Flp recombinase, Ten3 protein is tagged with Halo. Bottom: Ten3 protein expression in a horizontal section across the hippocampal-entorhinal region in a postnatal day 8 (P8) mouse was visualized with anti-HA antibody in the hippocampus of *Ten3*$^{cTAG/cTAG}$ mice (left), and with Halo tag ligand in the same area in *Pgk1-FlpO;Ten3*$^{cTAG/cTAG}$ mice. Both samples are double stained with HA and Halo tag ligand, demonstrating lack of Halo signal in the absence of Flp and complete conversion of HA into Halo in the presence of Flp.

(B) Same as (A) except for the *Lphn2*$^{cTAG}$ mice. In the absence of Flp recombinase, Lphn2 is tagged with a FLAG tag; in the presence of Flp recombinase, Lphn2 is tagged with a SNAP tag.

Scale bar, 200 µm.

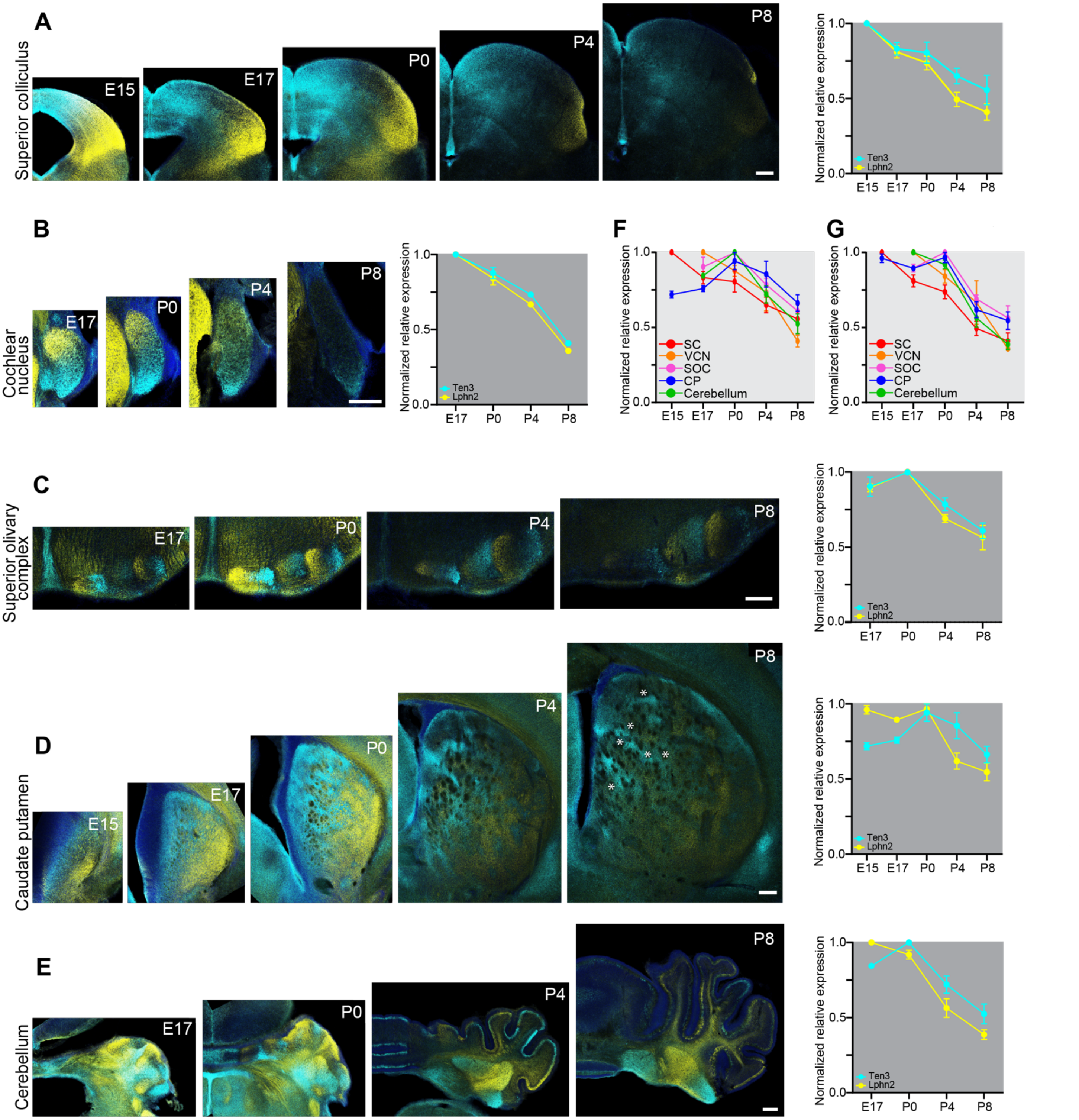


**Figure S2. Temporal patterns of Ten3 and Lphn2 expression across developing brain regions, related to Figures 2, 3, 4 and 6**

Coronal sections across developmental stages were stained in the same batch of experiments under identical conditions and imaged under identical exposure settings to allow for comparative analysis of relative protein intensity across time. Quantifications of normalized relative levels of expression are shown in the rightmost panels. All quantification analyses were based on n = 3 mice, except cochlear nucleus at P4 and P8 (n = 2). Scale bar = 200 µm for all images.

(A) Ten3 and Lphn2 protein expression in the superior colliculus (SC). Expression of both proteins peaked during embryogenesis at E15 or earlier, followed by a continuous decline after birth. The timing of peak expression coincided with the period when retinal ganglion cell (RGC) axons innervate the SC at E15,[S1] and the subsequent decline of expression corresponded to the onset of refinement of axon targeting after birth by cholinergic retinal waves.[S2]

(B, C) Ten3 and Lphn2 protein expression in the ventral cochlear nucleus (B) and the nuclei of the superior olivary complex (C). In the ventral cochlear nucleus (VCN), peak expression levels of both Ten3 and Lphn2 were observed during embryogenesis at E17 or earlier, with a drastic decrease in expression after P4. The peak expression of Ten3 and Lphn2 in the superior olivary complex (SOC) was at around birth with a sharp reduction at P4. These expression profiles aligned with the known developmental timeline of the auditory brainstem. During embryogenesis, peak expression in the VCN coincides with the period of initial axonal outgrowth at E15.5.[S3-5] This is followed by peak expression at the target, the SOC, at P0, when axons mature into a calyx in the medial nucleus of the trapezoid body (MNTB)[S6] and prune in the lateral superior olive (LSO), nuclei of the SOC. Expression declined by P4, a time when spontaneous cochlear activity begins to drive further synaptic refinement in the VCN and the SOC.[S7-10] Notably, neurons of the VCN are born in a dorsal–ventral birth order gradient, whereas those in the SOC are born in a medial–lateral gradient.[S11-13] Within both VCN and SOC, Lphn2$^{+}$ regions corresponded to earlier born neurons, whereas Ten3$^{+}$ regions corresponded to later-born neurons. Throughout development, Ten3 and Lphn2 maintained spatially inverse expression patterns.

(D) Ten3 and Lphn2 protein expression in the caudate putamen (CP, also known as striatum). Ten3 was highly enriched in dorsomedial CP, whereas Lphn2 was predominantly expressed in the ventrolateral CP, with highest expression of Lphn2 at E15 and Ten3 at P0. The peak expression of Lphn2 and Ten3 coincided with the timing of neuronal birth, where the spiny projection neurons constituting 95% of the CP neurons are born in a ventrolateral (E14–18) to dorsomedial (E18–P0) birth order gradient throughout the entire CP.[S14] Here again, Lphn2$^{+}$ regions corresponded to earlier born neurons, whereas Ten3$^{+}$ regions corresponded to later-born neurons. Also, earlier in development, the intermediate zone where Ten3 and Lphn2 domains converged appeared relatively homogeneous. By P0, these regions resolved into discrete Ten3- and Lphn2-enriched clusters, forming spatially refined hotspots. This transition paralleled the onset of glutamatergic innervation from both the thalamus[S15] and cortex,[S16,17] the maturation of synaptic connections, and dendritic spine growth in spiny projection neurons, during which bundles of corticostriatal axonal fibers begin to invade and penetrate the CP. * indicate example cortical axon bundles passing through the CP, creating "holes" in the Ten3 and Lphn2 expression patterns.

(E) Ten3 and Lphn2 protein expression in the cerebellum. Inverse expression of Ten3 and Lphn2 was observed in both Purkinje cells (PCs) and the cerebellar nuclei (CN) throughout the cerebellum. Peak expression of Lphn2 was observed E17, whereas peak Ten3 expression was at P0. As lobulation progressed postnatally, the emergence of stable, inversely patterned expression across the cerebellar cortex and nuclei along the medial–lateral axis was observed.

(F, G) Normalized relative Ten3 (F) and Lphn2 (G) protein expression in all areas throughout development.

Abbreviations: SC, superior colliculus; SOC, superior oliveary complex; VCN, ventral cochlear nucleus; CP, caudate putamen.

Together, these findings reveal that Ten3 and Lphn2 exhibit inverse expression that are both spatially segregated and temporally dynamic across multiple brain systems. Peak Ten3 and Lphn2 expression was observed during embryogenesis in the SC and cochlear nucleus, whereas expression in the SOC, CP, and cerebellum peaked around birth. Notably, in the auditory, striatal, and cerebellar circuits, Lphn2 expression correlated with earlier-born and Ten3 with later-born neurons, suggesting Ten3 and Lphn2 may establish connectivity across temporally distinct neuronal populations through sequential molecular expression. In systems where timing of target selection of axons has been determined, this timing overlaps with peak expression of Ten3 and Lphn2.

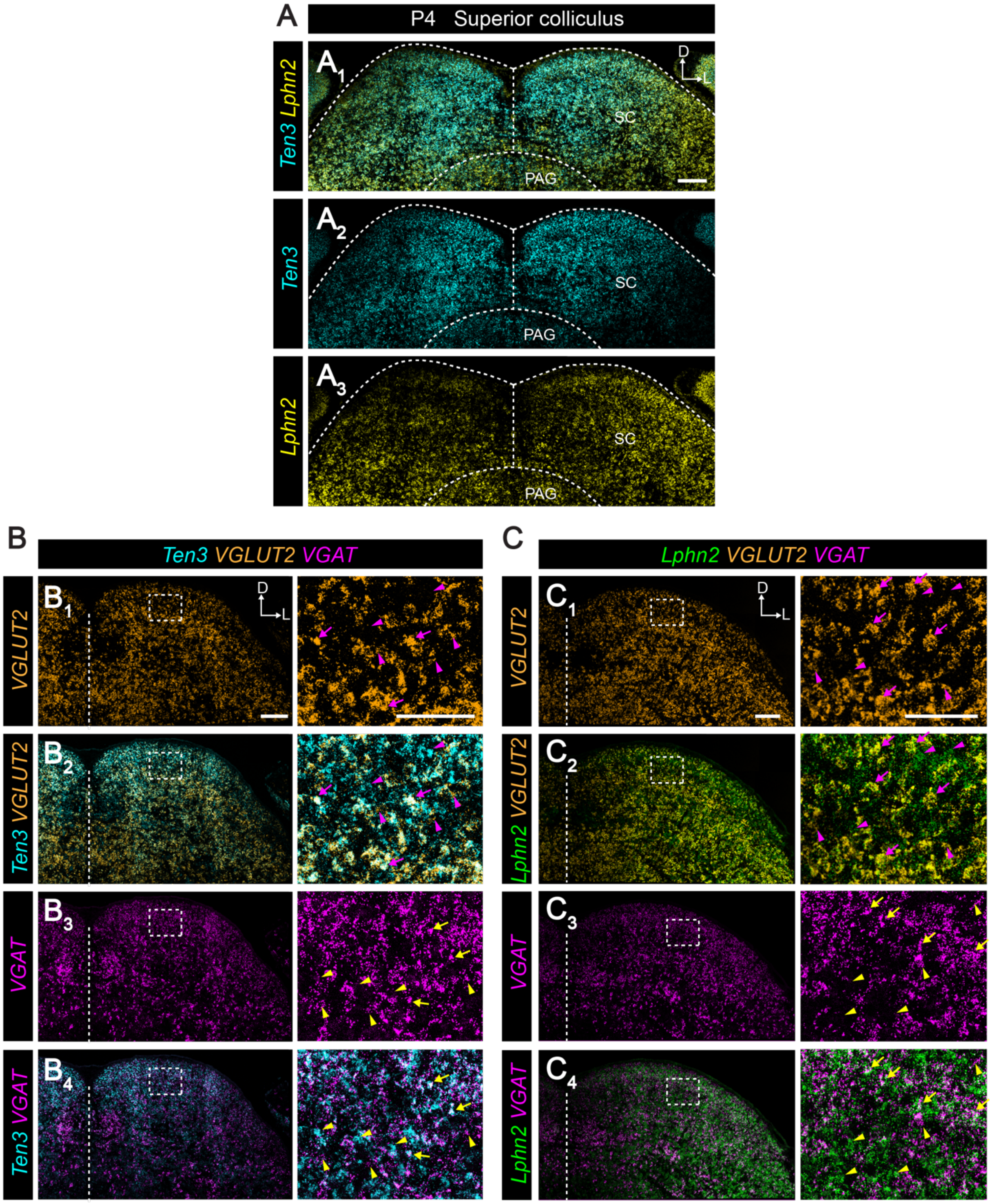


**Figure S3. *Ten3* and *Lphn2* mRNA display inverse expression in the superior colliculus, related to Figure 2**

(A) *Ten3* and *Lphn2* mRNA expression in a single coronal section of the superior colliculus at P4.

(B, C) Co-expression analysis of *Ten3* mRNA (B) and *Lphn2* mRNA (C) with *VGLUT2* and *VGAT* mRNA, markers for excitatory and inhibitory neurons, respectively. In the high-magnification insets (correspond to the dashed rectangles to the left), arrows point to cells co-expressing *Ten3* or *Lphn2* and *VGLUT2* or *VGAT*. Side arrowheads point to cells only expressing either *Ten3* mRNA (B) or *Lphn2* mRNA (C). Upward arrowheads point to cells only expressing either *VGLUT2* mRNA (top two panels) or *VGAT* mRNA (bottom two panels). Both *Ten3* and *Lphn2* are expressed in both excitatory and inhibitory neurons. Vertical dashed line indicates the midline. Scale bar, 200 μm (inset 100 μm).

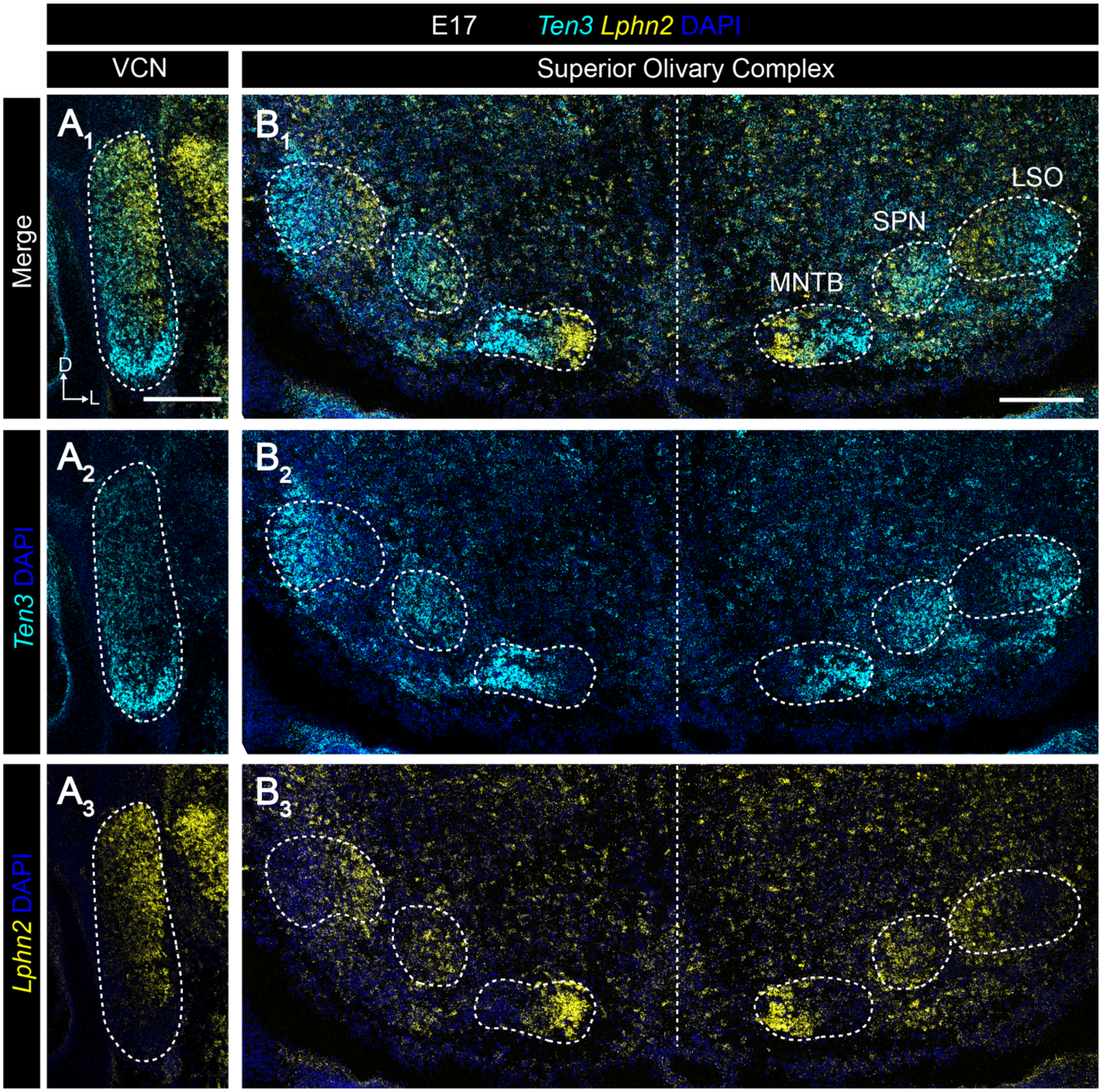


**Figure S4. *Ten3* and *Lphn2* mRNA display inverse expression in the auditory brainstem nuclei, related to Figure 3**

*Ten3* and *Lphn2* mRNA expression in a single coronal section of the ventral cochlear nucleus (VCN; A) and the nuclei of the superior olivary complex (B) at E17. VCN, MNTB, SPN, and LSO are demarcated by the dashed outlines. MNTB, medial nucleus of the trapezoid body; SPN, superior paraolivary nucleus; LSO, lateral superior olive. Vertical dashed line indicates the midline. Scale bar, 200 μm.

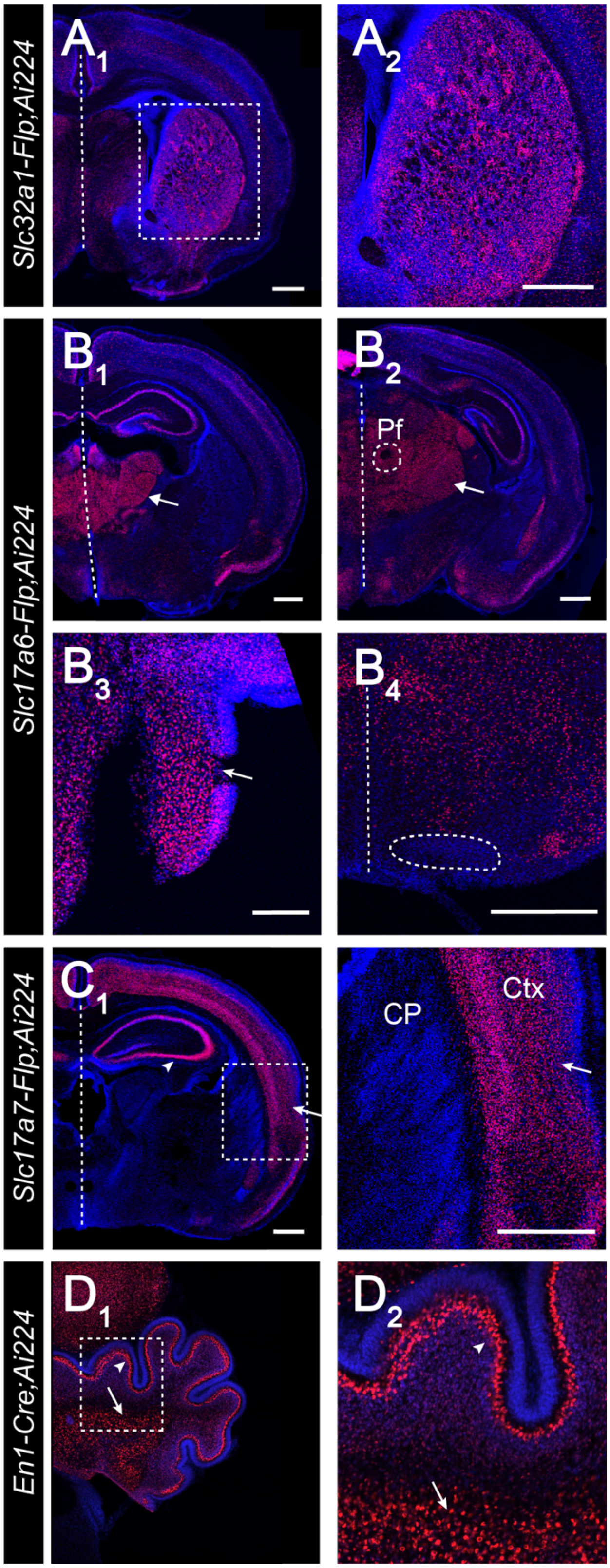


**Figure S5. Validation of Flp and Cre driver lines, related to Figures 3, 5, and 7**

(A) Coronal section of *Slc32a1-Flp;Ai224* at P4 showing labeled GABAergic neurons in the CP. *Ai224* is a Flp-dependent nuclear tdTomato reporter line. $A_2$ is a higher-magnified view of the white dashed box in $A_1$, highlighting labeled GABAergic neurons within CP. Vertical dashed lines are midlines in this and all other panels.

(B) Coronal sections of *Slc17a6-Flp;Ai224* at P4 showing labeled excitatory neurons in the anterior thalamus ($B_1$, arrow), posterior thalamus ($B_2$, arrow) including parafascilular nucleus (Pf, demarcated by dashed circle in

$B_2$), and ventral cochlear nucleus ($B_3$, arrow). However, cells in the medial nucleus of the trapezoid body of the superior olivary complex demarcated by the dashed outlines are not labeled ($B_4$).
(C) Coronal section of *Slc17a7-Flp;Ai224* at P4 showing labeled excitatory neurons in the cortex. $C_2$ is a higher-magnified view of the white dashed box in $C_1$. Arrowhead and arrow point to labeled excitatory neurons within the hippocampus and the cortex, respectively.
(D) Coronal section of *En1-Cre;Ai224* at P4 showing labeled Purkinje cells and the neurons in the cerebellar nuclei. $D_2$ is a higher-magnified view of the white dashed box in $D_1$, highlighting labeled neurons within the cerebellar nuclei (arrow) and the Purkinje cell layer (arrowhead).
CP, Caudate putamen; Ctx, Cortex. Dashed vertical line, midline. Scale bar, 500 µm (100 µm for $B_3$).

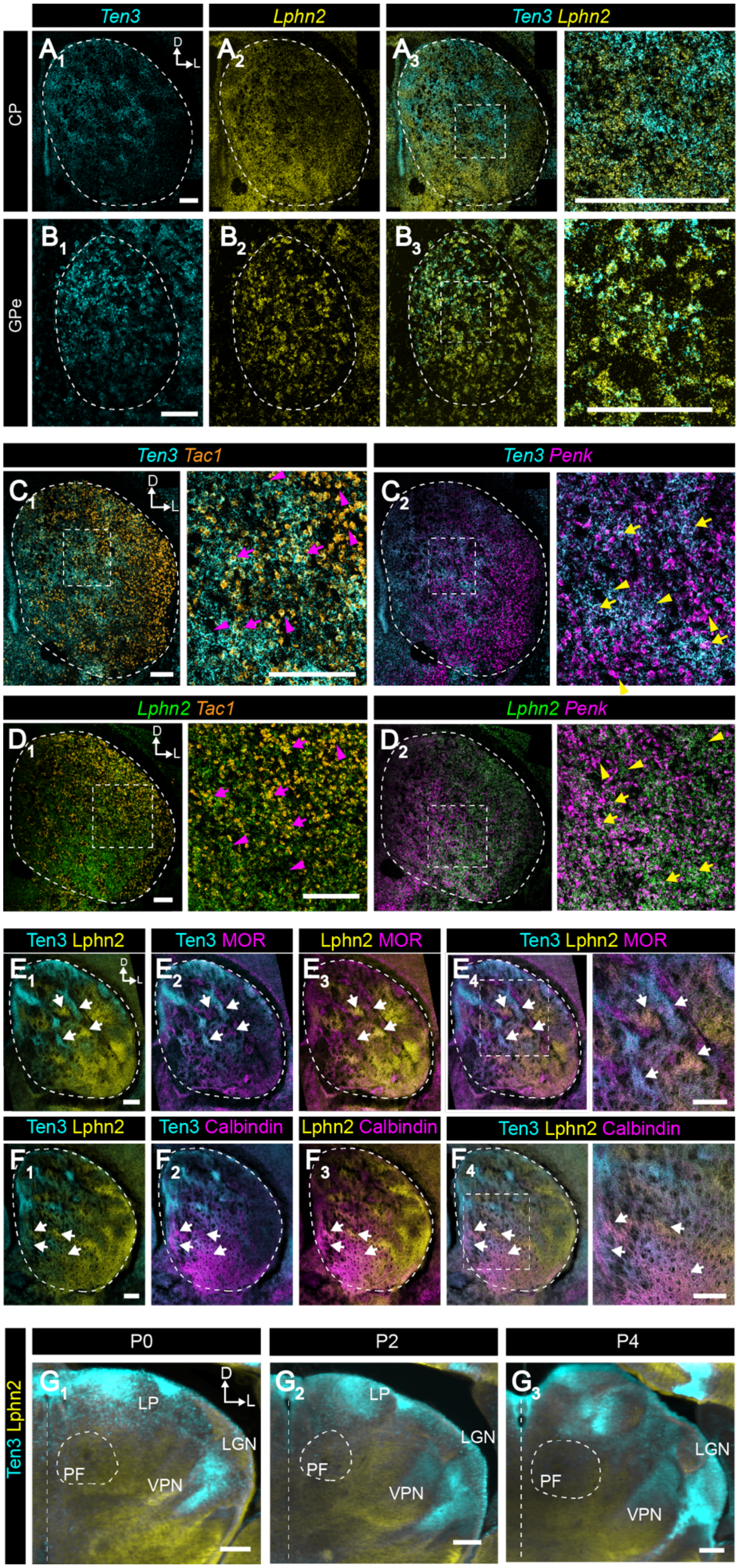


**Figure S6. *Ten3* and *Lphn2* mRNA display inverse expression in CP and GPe, related to Figure 4**
(A, B) *Ten3* and *Lphn2* mRNA expression in the CP (A) and GPe (B) of a P4 brain, shown in coronal sections. CP and GPe are demarcated by the dashed outlines based on DAPI co-staining (not shown). High-magnification insets correspond to the dashed rectangles in the left panel.

(C, D) Co-expression analysis of *Ten3* (E) and *Lphn2* (F) with *Tac1* ($C_1$; $D_1$) or *Penk* ($C_2$; $D_2$), markers of direct and indirect pathway spiny projection neurons, respectively. High-magnification insets correspond to the dashed rectangles in the left panel. Arrows point to cells co-expressing *Ten3* (C) or *Lphn2* (D) and *Tac1*$^+$ or *Penk*$^+$ CP regions. Side arrowheads point to cells exclusively expressing either *Ten3* mRNA (C) or *Lphn2* mRNA (D). Upward arrowheads point to cells exclusively expressing either *Tac1* mRNA ($C_1$; $D_1$) or *Penk* mRNA ($C_2$; $D_2$). Some *Ten3*$^+$ regions overlap with *Tac1* expression and others overlap with *Penk* expression; likewise for *Lphn2*$^+$ regions. Thus, both *Ten3* and *Lphn2* are expressed in spiny projection neurons in the direct and indirect pathways.
(E, F) Co-expression analysis of Ten3 and Lphn2 with μ-opioid receptor (MOR, E) or calbindin (F), markers of striosomes and extrastriosomal matrix, respectively. High-magnification insets correspond to the dashed rectangles in the left panel ($E_4$; $F_4$). Arrows point to cells co-expressing Ten3 or Lphn2 and MOR$^+$ (E) or Calbindin$^+$ (F) CP regions. As is evident, some Ten3$^+$ hotspots overlap with MOR expression and others overlap with calbindin expression; likewise for Lphn2$^+$ hotspots. Thus, Ten3 and Lphn2 hotspots do not correspond with the striosome/matrix division of the CP.
(G) Ten3 and Lphn2 protein expression in the thalamus at P0, 2 and 4. Note that parafascicular nucleus, a nucleus known to project axons to the CP, does not express high levels of Ten3 protein during development, while other thalamic nuclei, including the lateral posterior nucleus of the thalamus (LP), lateral geniculate nucleus (LGN), and ventral posterior nucleus of the thalamus (VPN), display high Ten3 expression.
Scale bar, 200 μm.

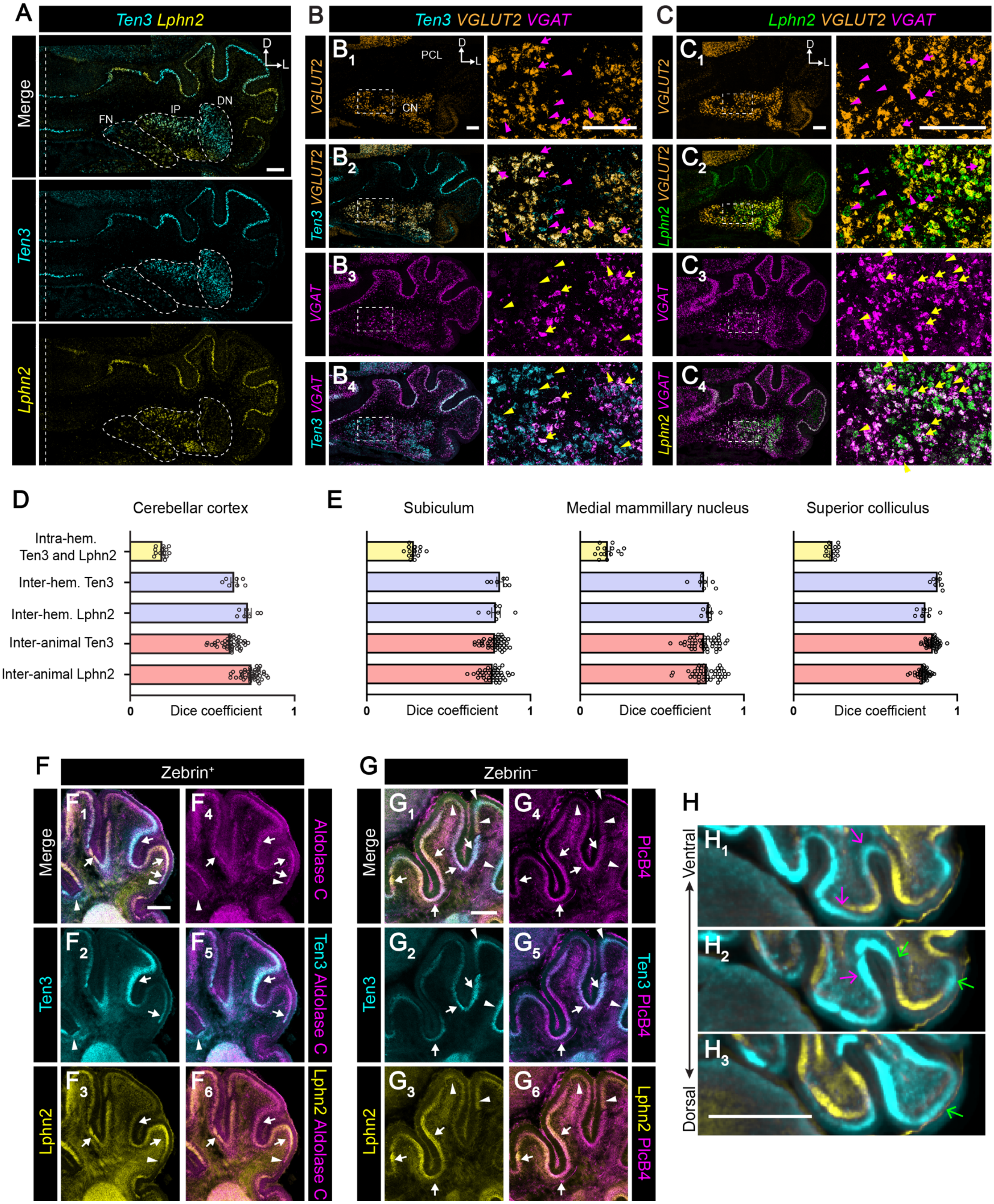


**Figure S7. *Ten3* and *Lphn2* mRNA and proteins co-expression analysis in the cerebellum, related to Figure 6**

(A) *Ten3* and *Lphn2* mRNA expression in the cerebellum of a P4 brain, shown in a coronal section, showing their inverse expression along the medial–lateral axis. Notably, *Ten3* and *Lphn2* expression in the CN did not conform to the anatomical boundaries of the fastigial, interposed, and dentate nuclei of CN, demarcated by dashed outlines, suggesting their expression is not restricted to specific anatomical CN subdivisions but instead reveals a new anatomical organization. Vertical dashed line indicates the midline. FN, fastigial nucleus; IP, interposed nucleus; DN, dentate nucleus.

(B, C) Co-expression of *Ten3* mRNA (C) or *Lphn2* mRNA (D) with both *VGLUT2* or *VGAT* mRNA, markers for excitatory or inhibitory neurons, respectively, in the cerebellar nuclei. In high-magnification insets, arrows point to cells that co-express *Ten3* or *Lphn2* with either *VGLUT2* or *VGAT*. Side arrowheads point to cells only expressing either *Ten3* mRNA (C) or *Lphn2* mRNA (D). Upward arrowheads point to cells only expressing either *VGLUT2* mRNA (top two panels) or *VGAT* mRNA (bottom two panels). These data indicate that both *Ten3* and *Lphn2* are expressed in both glutamatergic and GABAergic neurons in the cerebellar nuclei. PCL, Purkinje cell layer; CN, cerebellar nuclei.

(D) Quantification of spatial stereotypy of Ten3 and Lphn2 expression in 3D P0 cerebella (n = 7 brains) using the Dice similarity coefficient (DSC). DSC measures the accuracy of spatial overlap, where 0 represents complete spatial segregation and 1 represents complete spatial overlap. After signal segmentation and whole brain registration (see STAR Methods for detail), DSC analysis revealed minimal intra-hemispheric overlap between Ten3 and Lphn2 (top row)—consistent with their inverse expression patterns, and high inter-hemispheric ($2^{nd}$ and $3^{rd}$ rows) and inter-animal similarity ($4^{th}$ and $5^{th}$ rows) for either Ten3 or for Lphn2—indicating robust spatial stereotypy. Each dot represents a pairwise comparison; error bars represent mean ± SEM.

(E) As a comparision of spatial stereotypy of Ten3 and Lphn2 expression in the cerebellum (D), here we quantify spatial stereotypy of Ten3 and Lphn2 expression in the P0 subiculum, medial mammillary nucleus, and the superior colliculus using DSC. Comparisons include spatial overlap of Ten3 and Lphn2 within the same hemisphere (top), Ten3 or Lphn2 expression across hemispheres of the same brain ($2^{nd}$ and $3^{rd}$ rows), and inter-animal comparisons of Ten3 and Lphn2 expression patterns ($4^{th}$ and $5^{th}$ rows). Each dot represents a pairwise comparison; bar represent mean ± SEM.

(F, G) Ten3 and Lphn2 protein expression in the cerebellum of a P4 brain partially overlap with both aldolase C (F) and PlcB4 (G), zebrin$^{+}$ and zebrin$^{-}$ markers, respectively. Arrows indicate regions where either Ten3 or Lphn2 expression overlaps with zebrin$^{+}$ (F) or zebrin$^{-}$ (G) markers, and arrowheads indicate where either Ten3 or Lphn2 expression does not overlap with zebrin$^{+}$ (F) or zebrin$^{-}$ (G) markers. A subset of Aldolase C$^{+}$ regions overlapped with either Ten3$^{+}$ and Lphn2$^{+}$ domains, and similarly, a subset of Plcb4$^{+}$ regions comprised regions overlapping with either Ten3 or Lphn2 expression. These data indicate that both Ten3 and Lphn2 are expressed in both zebrin$^{+}$ and zebrin$^{-}$ neurons.

(H) Ten3 and Lphn2 protein expression in the cerebellar cortex visualized in serial horizontal sections following iDISCO clearing and whole brain light-sheet imaging. Magenta arrows in $G_1$ indicate two discrete Ten3-expressing domains that converge in more dorsal plane ($H_2$). Green arrows in $G_2$ mark another pair of Ten3 domains that merge further dorsally ($G_3$). In 3D, both Ten3 and Lphn2 expression form continuous, spatially distinct domains, producing five discrete domains per hemisphere. Scale bar, 700 µm.

Scale bar, 200 µm for all panels, unless specified.
